# Structures of human antibodies bound to SARS-CoV-2 spike reveal common epitopes and recurrent features of antibodies

**DOI:** 10.1101/2020.05.28.121533

**Authors:** Christopher O. Barnes, Anthony P. West, Kathryn E. Huey-Tubman, Magnus A.G. Hoffmann, Naima G. Sharaf, Pauline R. Hoffman, Nicholas Koranda, Harry B. Gristick, Christian Gaebler, Frauke Muecksch, Julio C. Cetrulo Lorenzi, Shlomo Finkin, Thomas Hagglof, Arlene Hurley, Katrina G. Millard, Yiska Weisblum, Fabian Schmidt, Theodora Hatziioannou, Paul D. Bieniasz, Marina Caskey, Davide F. Robbiani, Michel C. Nussenzweig, Pamela J. Bjorkman

## Abstract

Neutralizing antibody responses to coronaviruses focus on the trimeric spike, with most against the receptor-binding domain (RBD). Here we characterized polyclonal IgGs and Fabs from COVID-19 convalescent individuals for recognition of coronavirus spikes. Plasma IgGs differed in their degree of focus on RBD epitopes, recognition of SARS-CoV, MERS-CoV, and mild coronaviruses, and how avidity effects contributed to increased binding/neutralization of IgGs over Fabs. Electron microscopy reconstructions of polyclonal plasma Fab-spike complexes showed recognition of both S1^A^ and RBD epitopes. A 3.4Å cryo-EM structure of a neutralizing monoclonal Fab-S complex revealed an epitope that blocks ACE2 receptor-binding on “up” RBDs. Modeling suggested that IgGs targeting these sites have different potentials for inter-spike crosslinking on viruses and would not be greatly affected by identified SARS-CoV-2 spike mutations. These studies structurally define a recurrent anti-SARS-CoV-2 antibody class derived from *VH3-53/VH3-66* and similarity to a SARS-CoV *VH3-30* antibody, providing criteria for evaluating vaccine-elicited antibodies.

## Introduction

A newly-emergent betacoronavirus, SARS-CoV-2, resulted in a pandemic in 2020, causing the respiratory disease COVID-19 (Wu et al., 2020b; Zhou et al., 2020). SARS-CoV-2 is the third zoonotic betacoronavirus to infect humans this century, following SARS-CoV and MERS-CoV (Middle East Respiratory Syndrome) infections in 2003 and 2012, respectively (de Wit et al., 2016). In addition, four globally-distributed human coronaviruses, HCoV-OC43, HCoV-HKU1 (betacoronaviruses), and HCoV-NL63, HCoV-229E (alphacoronaviruses), contribute to 15-30% of common colds (Fung and Liu, 2019).The neutralizing antibody response to coronaviruses is primarily directed against the trimeric spike glycoprotein (S) on the viral membrane envelope, which serves as the machinery to fuse the viral and host cell membranes (Fung and Liu, 2019). Coronavirus S proteins contain three copies of an S1 subunit comprising the S1^A^ through S1^D^ domains, which mediates attachment to target cells, and three copies of an S2 subunit, which contains the fusion peptide and functions in membrane fusion (Figure 1A). Neutralizing antibody responses against SARS-CoV-2, SARS-CoV, and MERS-CoV S proteins often target the receptor-binding domain (RBD; also called the S1^B^ domain) (Hwang et al., 2006; Pinto et al., 2020; Prabakaran et al., 2006; Reguera et al., 2012; Rockx et al., 2008; Walls et al., 2020; Walls et al., 2019; Widjaja et al., 2019; Wrapp and McLellan, 2019; Wrapp et al., 2020).

**Figure 1.**
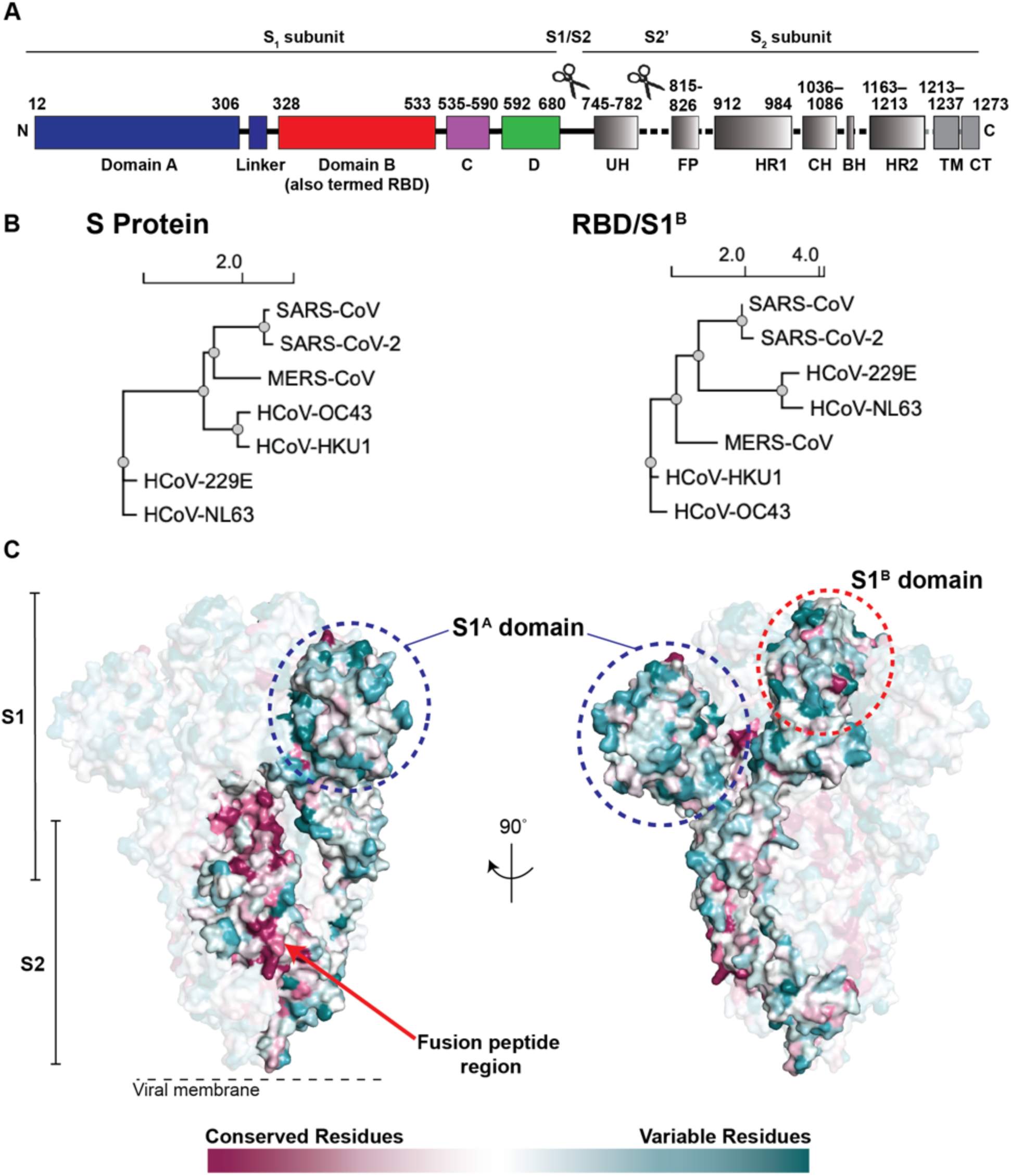
Coronavirus S proteins show localized regions of conservation and variability. (A) Schematic of SARS-CoV-2 S protein domain architecture. The S1 and S2 subunits are indicated, with scissors representing the locations of proteolytic cleavage sites required for S priming prior to fusion. UH = upstream helix, FP = fusion peptide, HR1 = heptad repeat 1, CH = central helix, BH = β-hairpin, HR2 = heptad repeat 2, TM = transmembrane region, CT = cytoplasmic tail. (B) Phylogenetic trees of selected coronaviruses based on protein sequences of S proteins and RBD/S1^B^ domains. (C) Sequence conservation of 7 human coronaviruses plotted as a surface. The sequence alignment was generated using SARS-CoV-2 (GenBank MN985325.1), SARS-CoV (AAP13441.1), MERS-CoV (JX869059.2), HCoV-OC43 (AAT84362.1), HCoV-229E (AAK32191.1), HCoV-NL63 (AAS58177.1), and HCoV-HKU1 (Q0ZME7.1). Conservation was calculated by ConSurf Database (Landau et al., 2005) and displayed using a surface representation of the structure of the SARS-CoV-2 S protein (PDB code 6VXX).

The S proteins of SARS-CoV-2 (1273 residues, strain Wuhan-Hu-1) and SARS-CoV (1255 residues, strain Urbani) share 77.5% amino acid sequence identity, while the S proteins of SARS-CoV-2 and MERS-CoV (1353 residues, strain EMC2012) are more distantly related, sharing only 31% identity (Figure 1B,C). Sequence identities between SARS-CoV-2 and common cold coronavirus S proteins are even lower, varying between 25% and 30%. Phylogenetic analyses confirm that SARS-CoV-2 and SARS-CoV are more closely related to each other than to other human coronaviruses (Figure 1B). The RBD/S1^B^ domains show varying degrees of sequence identity, ranging from 13% (SARS-CoV-2 and HCoV-NL63) to 74% (SARS-CoV-2 and SARS-CoV). Nevertheless, the 3D structures of S protein trimer ectodomains are similar to each other and to other coronavirus S structures, including the finding of flexible RBDs (S1^B^ domains) that can be in various “up” conformations or in the “down” conformation of the closed pre-fusion trimer (Kirchdoerfer et al., 2016; Li et al., 2019; Walls et al., 2020; Walls et al., 2016; Wrapp et al., 2020; Yuan et al., 2017). Primary amino acid sequence differences in the RBDs of SARS-CoV-2 and SARS-CoV compared with MERS-CoV (Figure 1B,C) result in binding to different host receptors: angiotensin-converting enzyme 2 (ACE2) for SARS-CoV-2 and SARS-CoV (Hoffmann et al., 2020; Li et al., 2003; Zhou et al., 2020) and dipeptidyl peptidase 4 for MERS-CoV (Raj et al., 2013). One of the common cold coronaviruses, HCoV-NL63, also uses its RBD (S1^B^) to bind ACE2, although its interactions differ structurally from RBD-ACE2 interactions of SARS-CoV-2 and SARS-CoV (Tortorici and Veesler, 2019), whereas HCoV-OC43 and HCoV-HKU1 uses their S1^A^ domains to bind host receptors including 9-*O*-acetylated sialic acids (Tortorici et al., 2019).

Understanding the antibody response to SARS-CoV-2 S protein is of critical importance because correlates of protection for vaccines usually involve antibodies (Plotkin, 2001, 2008, 2010). Moreover, antibodies are being considered as therapeutics for COVID-19 patients (Zhou and Zhao, 2020). Relatively little is known about antibody recognition of SARS-CoV-2 S compared with other coronavirus S proteins (Graham et al., 2013; Gralinski and Baric, 2015; Wan et al., 2020). However, structures of S trimer, RBD-Fab, RBD-ACE2, and S trimer-Fab complexes for SARS-CoV-2 and other coronaviruses are informative for interpreting and understanding the antibody response to SARS-CoV-2 (Gui et al., 2017; Kirchdoerfer et al., 2020; Kirchdoerfer et al., 2016; Kirchdoerfer et al., 2018; Pallesen et al., 2017; Pinto et al., 2020; Shang et al., 2020; Shang et al., 2018; Walls et al., 2016; Walls et al., 2017; Walls et al., 2019; Wang et al., 2020; Xiong et al., 2018; Yuan et al., 2020).

Here, we analyzed purified IgG and Fabs from the plasmas of 10 COVID-19 convalescent individuals (Robbiani et al., 2020) for binding to trimeric S and monomeric RBD/S1^B^ domains of six human coronaviruses and for neutralization of SARS-CoV-2 pseudoviruses. To better understand the binding mechanism of polyclonal antibodies, we further characterized plasma Fabs from two individuals using negative-stain electron microscopy polyclonal epitope mapping (nsEMPEM), showing that the polyclonal landscape includes antibodies that target epitopes in both SARS-CoV-2 S1^A^ and RBD domains. In addition, we solved a 3.4 Å single-particle cryo-EM structure of an S trimer bound to a neutralizing monoclonal antibody (mAb), which targeted an epitope on an “up” RBD that overlapped with the RBD epitope identified by nsEMPEM and would sterically block ACE2 receptor binding. The epitopes we found represent binding classes defined by distinct VH gene segments, suggesting that these recurring classes are commonly represented in neutralizing antibodies against SARS-CoV-2 and provide criteria for evaluating neutralizing antibodies raised by infection or vaccination. Finally, we used modeling to suggest that distinct binding orientations allow for differential avidity effects, demonstrating the potential for inter-spike crosslinking that would increase effective affinities for some anti-S IgGs on SARS-CoV-2 virions.

## Results

### Convalescent plasma IgG and Fab binding properties demonstrate recognition of diverse coronaviruses and effects of avidity

Convalescent plasma samples were collected from individuals who had recovered from COVID-19 at Rockefeller University Hospital (Robbiani et al., 2020). We isolated polyclonal IgGs from 10 convalescent plasmas (Figure 2), most of which had high neutralizing titers (Robbiani et al., 2020), and compared binding of their IgGs to purified S proteins from SARS-CoV-2, SARS-CoV, MERS-CoV, HCoV-OC43, HCoV-NL63, and HCoV-229E (Figure S1) by ELISA (Figure 3; Figure S2). Purified plasma IgGs recognized S proteins from all coronaviruses evaluated, with weaker binding observed for most samples to MERS-CoV (Figure 3C) and common cold coronavirus S proteins (Figure 3D-F). Amongst the plasmas (COV21, COV57, and COV107) chosen for further analysis based on ELISA EC_50_ values and strong neutralization potencies (Robbiani et al., 2020), IgGs from COV21 and COV57 showed the strongest binding to the S proteins from SARS-CoV-2 and SARS-CoV, with only the COV57 IgGs showing measurable binding to MERS-CoV S protein. The COV107 IgGs showed intermediate binding to SARS-CoV-2 and SARS-CoV and no binding to MERS-CoV S proteins (Figure 3A-C).

**Figure 2.**
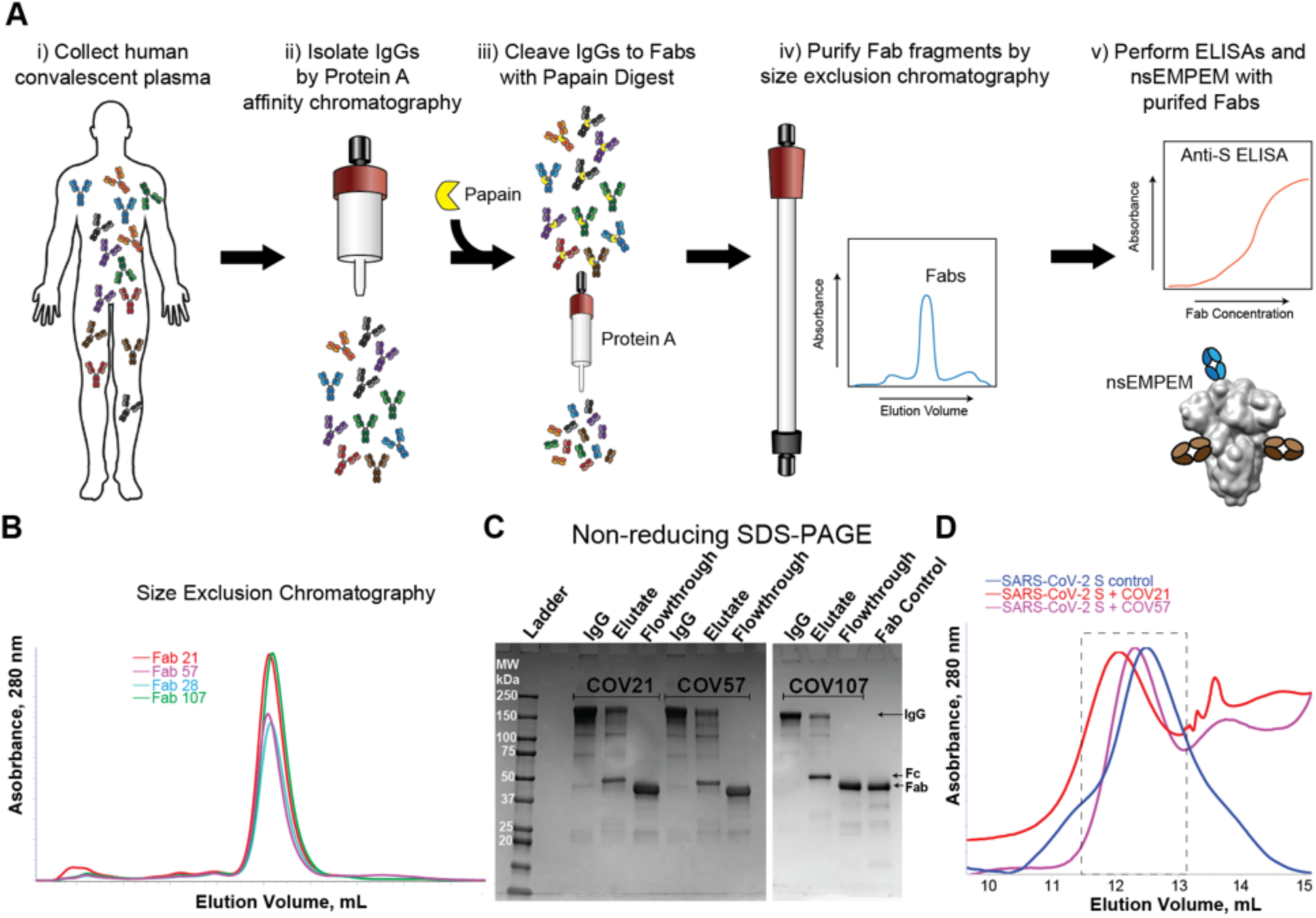
Plasma Fabs bind to SARS-CoV-2 S protein. (A) Schematic of polyclonal IgG and Fab purification from human plasma for nsEMPEM protocol. (B) SEC profile of Fabs (left) and SDS-PAGE of purified IgGs and Fabs (right) from COV21, COV57, and COV107 plasma samples. (C) SEC demonstration that plasma-derived Fabs from COV21 and COV57 shift the SARS-CoV-2 S protein trimer to a higher apparent molecular weight. No shift was observed when Fabs from COV107 were analyzed by SEC with S protein (data not shown). Fractions pooled and concentrated for nsEMPEM are boxed. See also Figure S1.

**Figure 3.**
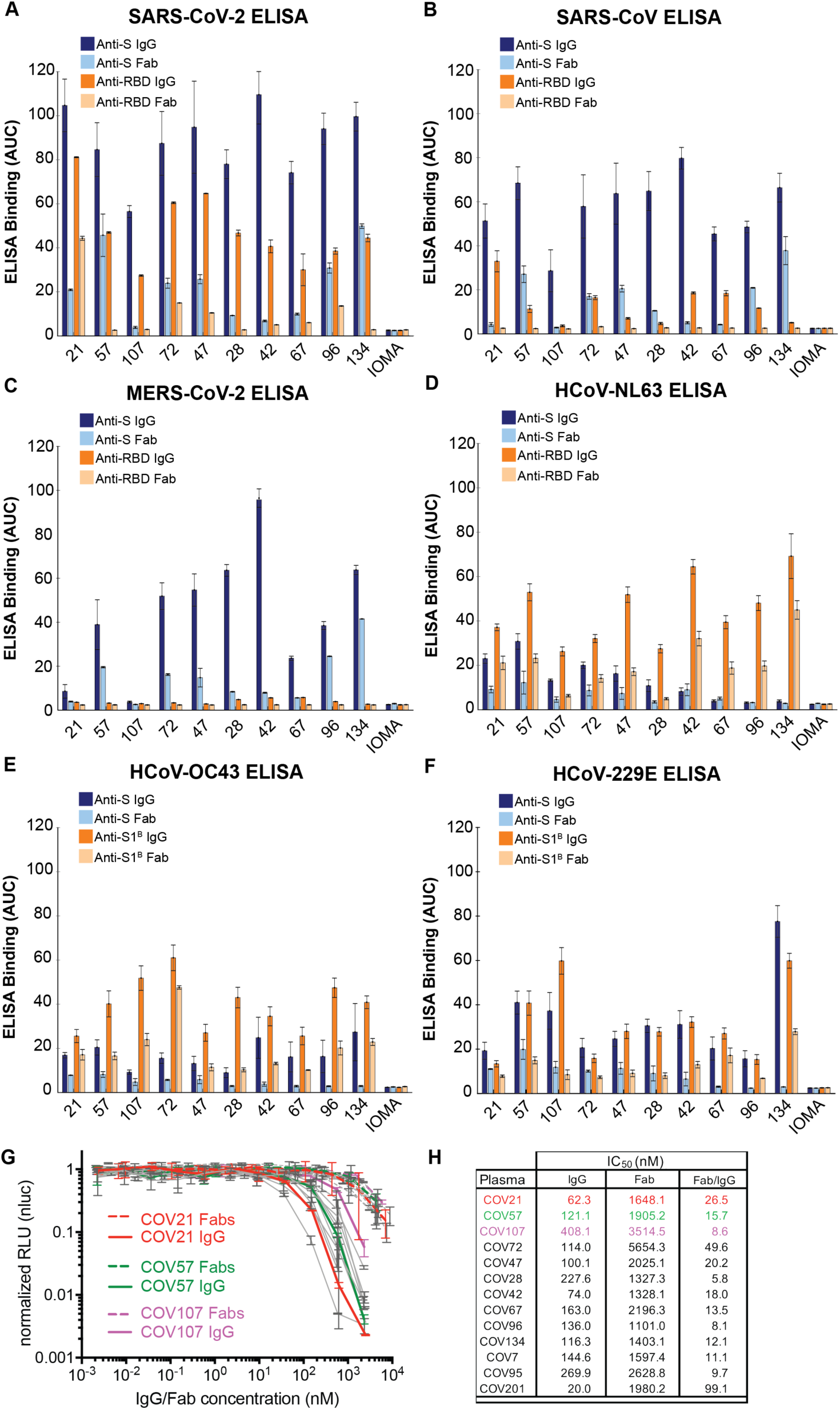
Convalescent plasma IgG and Fab binding properties demonstrate recognition of diverse coronaviruses and effects of avidity. Results from ELISAs assessing binding of IgGs and Fabs purified from plasmas from 10 COVID-19 individuals (X-axis) presented as area under the curve (AUC; shown as mean ± S.D. of values derived from experiments conducted in triplicate). Binding was assessed against S and RBD proteins for SARS-CoV-2 (A), SARS-CoV (B), MERS-CoV (C), HCoV-NL63 (D), HCoV-OC43 (E), and HCoV-229E (F). Polyclonal IgGs or Fabs were evaluated at a top concentration of 50 µg/mL and 7 additional 4-fold serial dilutions. Binding of the IgG and Fab from IOMA, an antibody against HIV-1 (Gristick et al., 2016), was used as a control in each assay. (G) In vitro neutralization assays comparing the potencies of purified plasma IgGs and purified plasma Fabs. COV21, COV57, and COV107 plasma Fabs and IgGs are highlighted in the indicated colors; curves for 10 other plasmas (listed in panel H) are gray. (H) Molar IC_50_ values for purified plasma IgGs and Fabs for the indicated plasmas are listed with the molar ratio for IC_50_ (Fab) to IC_50_ (IgG) shown in the right column. See also Figure S2, S3.

ELISAs against RBD (or S1^B^ domain for two of the common cold coronavirus S proteins) showed the strongest binding to SARS-CoV-2 RBD for COV21, followed by COV57 and then COV107 IgGs, with the proportion of RBD versus S binding from COV21, COV72, and COV47 suggesting that the majority of the IgG responses from these plasmas were focused on the RBD, often a target of neutralizing antibodies in coronavirus infections (Hwang et al., 2006; Pinto et al., 2020; Prabakaran et al., 2006; Reguera et al., 2012; Rockx et al., 2008; Walls et al., 2020; Walls et al., 2019; Widjaja et al., 2019; Wrapp et al., 2020). The only appreciable reactivity with SARS-CoV and MERS-CoV RBDs was exhibited by COV21 IgG, which bound to SARS-CoV RBD (Figure 3B). Although we cannot determine whether the same IgGs are binding all three S proteins, the potential for cross-reactive binding of SARS-CoV antibodies was demonstrated for a mAb that was isolated from a SARS-infected individual, which was shown to recognize SARS-CoV and SARS-CoV-2 RBDs (Pinto et al., 2020). No reactivity with MERS-CoV RBD was observed for any of the polyclonal IgGs (Figure 3C). For most of the plasma IgGs, binding to the RBD was substantially weaker than binding to the counterpart S protein, with the exception of the strong COV21 and COV72 responses to the SARS-CoV-2 RBD. Most of the plasma IgGs exhibited stronger binding to the common cold S1^B^/RBDs than to the counterpart S protein trimers (Figure 3D-F).

To assess the degree to which cross-reactive recognition contributed to binding of plasma IgGs to RBD/S1^B^ domains, we repeated the ELISAs before and after adsorption with SARS-CoV-2 RBD-coupled resin or a control resin for five plasma IgG samples (Figure S3). As a positive control, purified IgGs incubated with the RBD resin showed little or no SARS-CoV-2 RBD binding (Figure S3A). Binding to SARS-CoV RBD was also reduced for the IgGs remaining after SARS-CoV-2 RBD adsorption (Figure S3B), suggesting cross-reactive recognition consistent with the 78% sequence conservation and structural homology of SARS-CoV-2 RBD and SARS-CoV RBD (Walls et al., 2020). By contrast, adsorption of plasma IgGs with SARS-CoV-2 RBD resins had only a modest effect on binding to common cold coronavirus RBDs (Figure S3D-F), consistent with little to no cross-reactive antibody recognition, likely due the low conservation between the SARS-CoV-2 RBD and mild coronavirus RBDs (Premkumar et al., 2020). We also note that IgGs from control plasmas collected from individuals not exposed to SARS-CoV-2 exhibited binding to common cold coronavirus RBDs that was not affected by SARS-CoV-2 RBD adsorption (Figure S3), again consistent with pre-exposure to mild coronaviruses rather than cross-reactivity with SARS-CoV-2 RBD.

Taken together, these results suggest: (i) The binding strengths and patterns of different coronavirus S protein recognition were diverse across COVID-19 individual plasma samples, (ii) Convalescent COVID-19 individuals harbor antibodies to the SARS-CoV-2 S protein, and to a lesser extent, the RBD/S1_B_, as well as reactivity to other coronaviruses, which likely represents previous exposure to common cold viruses, (iii) Polyclonal IgGs from individual plasma samples that bind to S proteins from MERS-CoV and/or SARS-CoV may display cross-reactive recognition, since the plasma donors were unlikely to have been infected with either of these coronaviruses, and (iv) Compared to the COV57 and COV107 plasmas, the COV21 IgG response had a higher proportion of IgGs that recognized the SARS-CoV-2 RBD.

We also evaluated the degree to which avidity effects contributed to the strength of binding of plasma IgGs to S proteins and RBDs by comparing the binding of bivalent polyclonal IgGs to monovalent Fabs, prepared by proteolytic cleavage of purified polyclonal IgGs (Figure 2B,C). Differential effects were evident in IgG to Fab comparisons: most of the SARS-CoV-2 anti-S response was reduced by at least 50% in the case of monovalent Fabs for all plasmas except for COV57 (Figure 3A). Recognition of the other coronavirus S proteins was also diminished for Fabs compared to intact IgGs (Figure 3B-F). For the three plasma IgGs that were further evaluated, the largest relative differences in IgG versus Fab binding to SARS-CoV-2 S protein was observed for COV21 and COV107; the IgG versus Fab binding difference for COV57 was less pronounced (Figure 3A). Notably, the SARS-CoV-2 S protein and RBD ELISAs showed that a higher fraction of the COV21 plasma IgGs were RBD-specific compared with the COV57 IgGs (Figure 3A,B) (Robbiani et al., 2020).

In summary, the ELISA data indicate that IgGs in plasma samples differ in their degree of focus upon epitopes within the S protein RBD/S1^B^ domain, their relative amounts of reactivity with SARS-CoV, MERS-CoV, and common cold coronaviruses, and the extent to which avidity effects contribute to the tighter binding of polyclonal bivalent IgGs as compared with monovalent Fabs.

### Plasma IgGs are more potent neutralizers than plasma Fabs

To investigate whether the bivalent architecture or larger size of IgGs compared with Fabs resulted in increased neutralization potencies, we measured the potencies of purified plasma IgGs and Fabs using in vitro neutralization assays (Figure 3G). SARS-CoV-2 pseudoviruses were constructed as described (Robbiani et al., 2020), and the concentrations of IgGs and Fabs at which 50% neutralization was achieved (IC_50_ values) were calculated. All tested plasma IgGs neutralized pseudoviruses at lower molar concentrations than their Fab counterparts, with increased potencies ranging from 6- to 100-fold (Figure 3H). The increased potency of the IgGs compared to Fabs was statistically significant (*p* = 0.0003), even when accounting for two Fabs per IgG. We conclude that bivalent IgGs more effectively neutralize SARS-CoV-2 pseudoviruses than monovalent Fabs.

### EM reveals distinct predominant epitopes targeted by convalescent plasma antibodies

We next used negative stain polyclonal electron microscopy (nsEMPEM) (Bianchi et al., 2018; Nogal et al., 2020) to map epitopes from Fabs isolated from convalescent COVID-19 plasma IgGs onto the SARS-CoV-2 S protein. In this method, Fabs that bind to an antigenic target are separated from non-binding Fabs in a polyclonal mixture by size-exclusion chromatography (SEC), Fab-antigen complexes are imaged by EM, and 2D/3D classification are used to identify predominant epitopes (Bianchi et al., 2018; Nogal et al., 2020) (Figure 2A-C). Typically, Fabs are incubated at 1000-2000x above EC_50_ values calculated from binding assays (Bianchi et al., 2018; Nogal et al., 2020). For most COVID-19 plasmas, Anti-S Fab EC_50_ values were estimated to be >50 µg/mL (Figure S2). However, purified polyclonal Fabs from COV21 and COV57 plasmas, which had approximate EC_50_s ranging from 20-50 µg/mL, showed stable binding by SEC after incubation with SARS-CoV-2 S trimers (Figure 2D), and 2D class averages showed evidence of bound Fabs (Figure S5). By contrast, purified Fabs from COV107 (EC_50_ >50 µg/mL) showed no evidence of binding to S by SEC (data not shown) or in a 3D reconstruction (Figure 4A).

**Figure 4.**
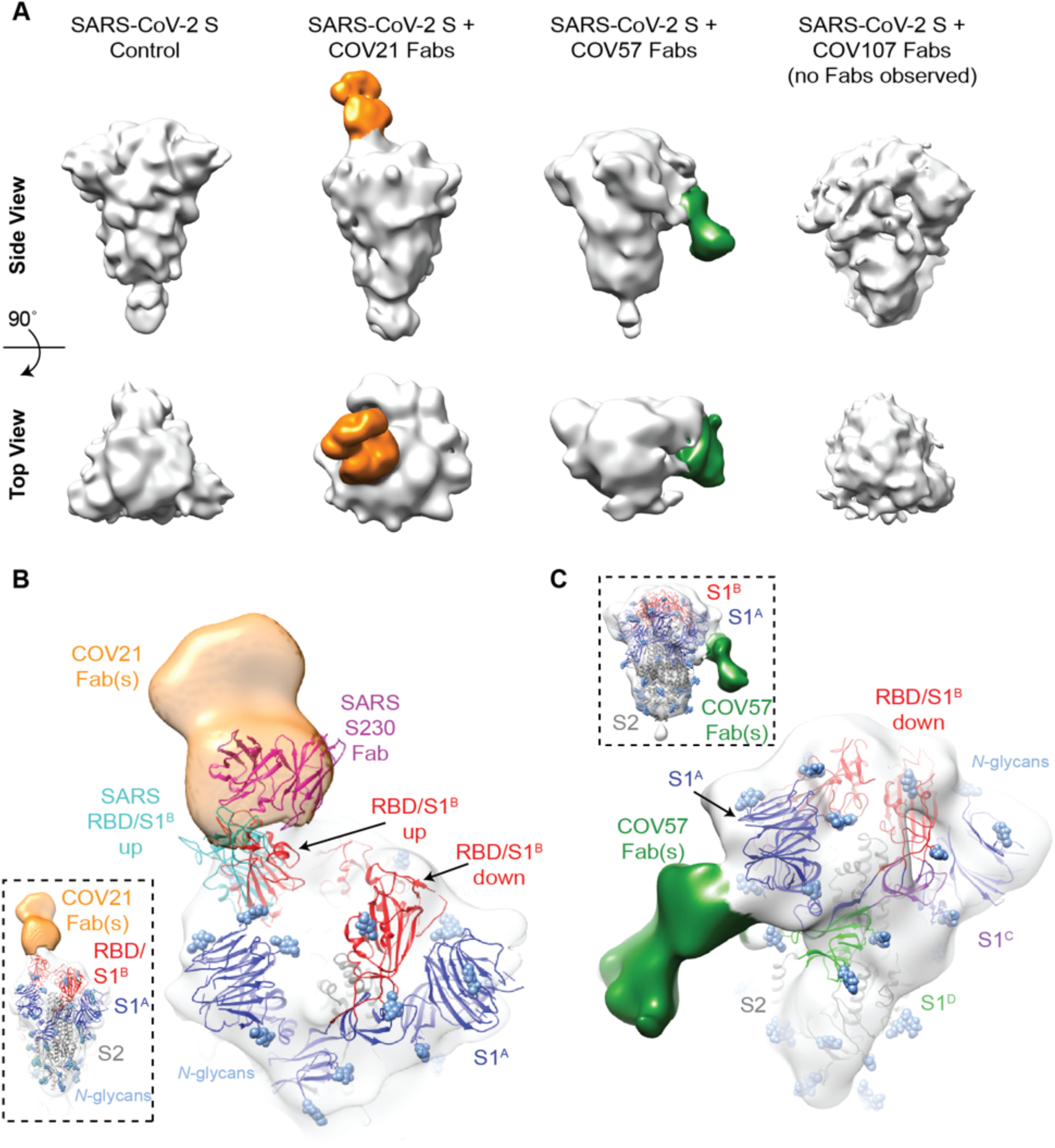
EM reveals distinct predominant epitopes targeted by convalescent plasma antibodies. (A) Side and top views for representative 3D reconstructions of four nsEMPEM datasets (S protein alone, S + COV21 Fabs, S + COV57 Fabs, S + COV107 Fabs). Bound Fabs observed in reconstructions from COV21 and COV57 plasmas are highlighted with false coloring as orange and green, respectively. No Fabs were observed in the reconstruction of COV107 Fabs plus S protein. Refined 3D models for SARS-CoV-2 S trimer-polyclonal Fab complexes from COV21 (panel B), and COV57 (panel C) were rigid-body fit with reference structures in Chimera (Goddard et al., 2007; Pettersen et al., 2004), displayed as cartoons (S1^A^: blue, S1^B^: red, S2: gray). (B) For COV21, the volume was best-fitted with PDB 6VYB (SARS-CoV-2, one “up” S1^B^ conformation, inset). Overlay of PDB 6NB6 showed similarities in S1^B^ epitope targeting of COV21 Fab (orange) and the human SARS-CoV neutralizing antibody, S230 (magenta, cartoon). (C) COV57 was fitted with PDB 6VXX (closed, prefusion conformation, inset). Fab density (green) was focused on the S1^A^ domain. See also Figure S4.

In order to verify that extra densities in nsEMPEM 3D reconstructions corresponded to bound Fab(s), we first solved a 3D reconstruction of SARS-CoV-2 S alone, revealing the expected low resolution structure of the closed, prefusion S trimer (Figure 4A). A 3D reconstruction of COV21 Fabs complexed with S showed recognizable density for the S trimer with a single extending density at the apex of the trimer corresponding to a Fab or mixture of Fabs bound to a similar epitope (Figure 4A). The density could be fit to an S trimer with a Fab bound to a single RBD in an “up” position using coordinates from SARS-CoV-2 S trimer structures (Walls et al., 2020; Wrapp et al., 2020), consistent with ELISA results mapping the COV21 response to the SARS-CoV-2 RBD (Figure 3A). The complex structure and the position of the COV21 Fab(s) closely resembled a structure of SARS-CoV S bound to a Fab from the S230 mAb isolated from a SARS-CoV–infected individual, whose epitope overlaps with the binding site for the ACE2 receptor (Walls et al., 2019) (Figure 4B). Interestingly, S230 binding was shown to functionally mimic ACE2 binding, allowing cleavage of the SARS-CoV S protein to promote fusogenic conformational rearrangements (Walls et al., 2019). While the COV21 Fab complex reconstruction showed occupancy for one S-protomer with an RBD in an “up” position (Figure 4A), COV21 Fab(s) could also bind analogous to the S230 Fab-SARS-Cov S complex, where classes of S trimer structures were found with two “up”/one “down” and three “up” RBD conformations (Walls et al., 2019).

Moreover, antibody S230, whose binding orientation resembles the position observed in the COV21 Fab(s) reconstruction (Figure 4B), appears to be a member of a class of recurrent anti-SARS mAbs. It belongs to a set of 10 non-clonally-related *VH3-30*-derived mAbs isolated from an individual infected with SARS-CoV, which represented 40% of the clones isolated from this individual (Pinto et al., 2020). Notably, these clones contained similar 9 amino acid CDRL3 sequences (consensus sequence MQGTHWPPT), suggesting that this group of mAbs has a common mode of binding, partially dependent on *VH3-30*-derived features. RBD residues 473 and 475 contacted by the antibody heavy chain in the S230 Fab–SARS-CoV structure (Walls et al., 2019) are conserved between SARS-CoV and SARS-CoV-2, and these residues are in the vicinity of antibody heavy chain residues N57 and K58. The only VH gene segments encoding the N57/K58 pair are *VH3-30, VH3-30-3*, and *VH3-33* (Lefranc et al., 2015). When mAbs were isolated after single B cell sorting using SARS-CoV-2 RBD as a bait, COV21 antibodies included heavy chains derived from *IGHV3-30*, which were also found in sequenced antibodies from five other donor plasmas (Robbiani et al., 2020). The similarity in binding orientation of COV21 Fab(s) with S230 (Figure 4B) suggests that COV21 Fab(s) may be members of the S230 recurrent class. Consistent with this hypothesis, 38 of 127 sequenced antibodies from the COV21 donor were derived from *VH3-30* or from the closely-related *VH3-30-3* or *VH3-33* VH gene segments (Robbiani et al., 2020).

The COV57 Fab(s)-S structure also showed recognizable density for both the S trimer and a single bound Fab(s) (Figure 4C). However, in this complex, the S trimer appeared closed with no RBDs in an “up” position, and the Fab density was not associated with an RBD, but rather with one of the S1^A^ subunits. In the complex, the Fab(s) pointed downwards (i.e., towards the viral membrane) rather that upwards (away from the viral membrane), as seen for the COV21 Fab(s). The COV57 Fab(s) density was in the vicinity of loops on the S1^A^ domain that were disordered in SARS-CoV-2 S trimer structures (Walls et al., 2020; Wrapp et al., 2020). Such flexibility could explain the diffuse nature of the COV57 Fab(s) density in this reconstruction. Interestingly, characterization of COV57 neutralization showed less correlation with RBD-specific antibodies relative to COV21 (Robbiani et al., 2020), consistent with the ELISAs (Figure 3A) and nsEMPEM characterizations (Figure 4C) reported here. This suggests that targeting S1 regions outside of the RBD may represent alternative modes for potent neutralization of SARS-CoV-2, as found for neutralizing antibodies isolated after vaccination against MERS-CoV in non-human primates (Wang et al., 2015).

### A cryo-EM structure of a monoclonal Fab-S protein complex resembles the COV21 Fab(s)-S reconstruction

Although we could not resolve densities for bound Fabs in the COV107-S nsEMPEM reconstruction (Figure 4A), RBD-binding mAbs isolated from the COV107 individual were potently neutralizing (Robbiani et al., 2020). We determined a 3.4 Å single-particle cryo-EM structure of the complex of one such antibody (C105; IC_50_ for neutralization of SARS-CoV-2 pseudovirus = 26.1 ng/mL) (Robbiani et al., 2020) bound to the SARS-CoV-2 S protein using a 1.8 Å crystal structure of the unbound C105 Fab for fitting to the cryo-EM density (Figure 5; Figures S5 S6, Tables S1,S2).

**Figure 5.**
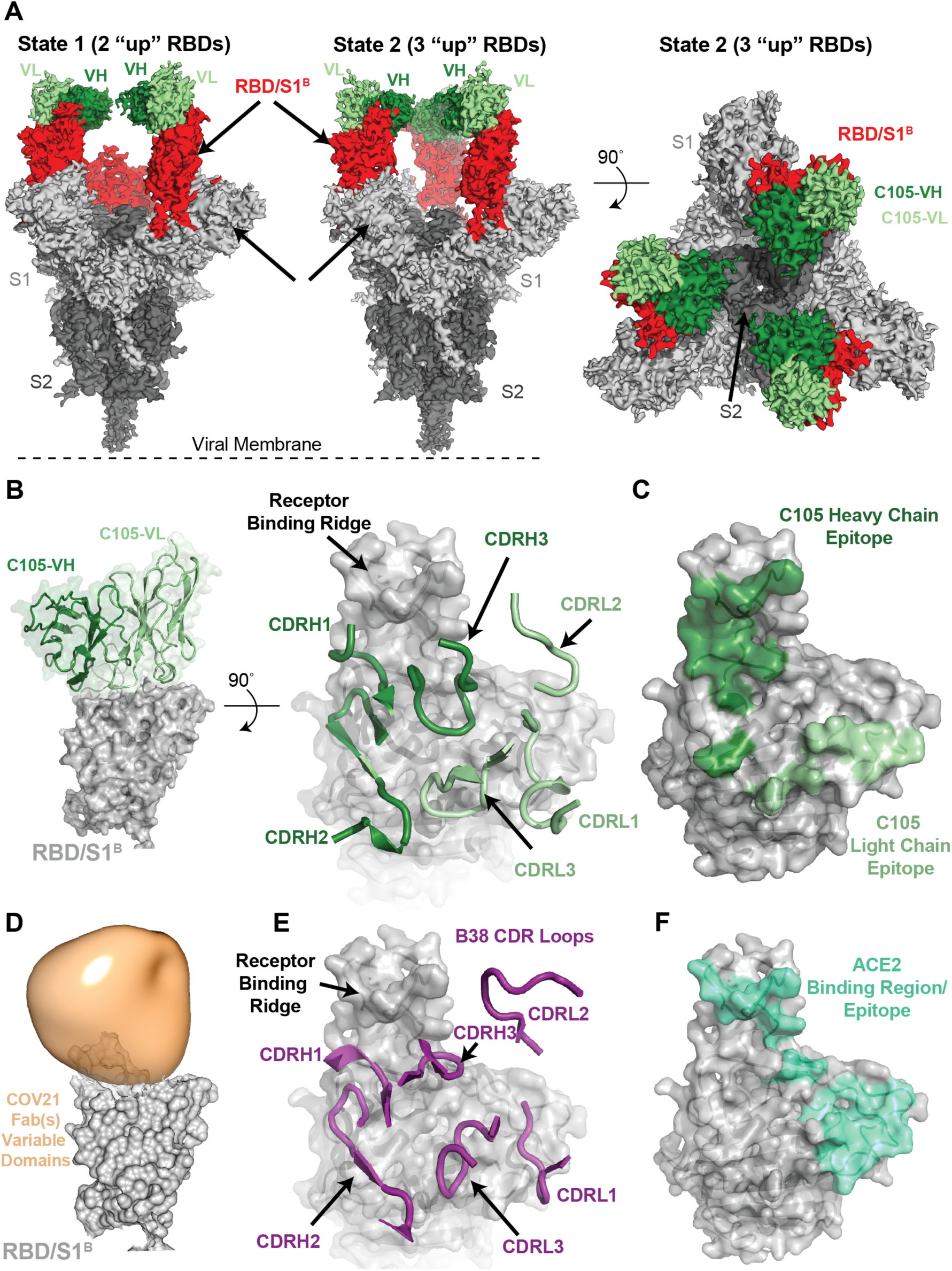
A cryo-EM structure of a monoclonal Fab-S protein complex resembles the COV21 Fab(s)-S reconstruction. (A) Reconstructed volumes for mAb C105 bound to SARS-CoV-2 S trimers in state 1 (two “up” RBDs, two bound Fabs) and state 2 (three “up” RBDs, three bound Fabs). (B) Left: Cartoon representation of VH-VL domains of C105 bound to an RBD. Right: CDR loops of C105 overlaid on surface representation of the RBD (shown as a gray surface). (C) RBD surface showing contacts by C105 VH-VL (contacts defined as an RBD residue within 7 Å of a VH or VL residue C*α* atom). (D) RBD surface fitted with volume representing the variable domains of the COV21 Fab(s) nsEMPEM reconstruction. (E) CDR loops of B38 mAb overlaid on surface representation of the RBD (from PDB code 7BZ5). (F) RBD surface showing contacts by ACE2 (contacts defined as an RBD residue within 7 Å of an ACE2 residue C*α* atom) (from PDB code 6VW1). See also Figure S5, S6.

We found two populations of C105 Fab-S complexes: an asymmetric S trimer with two “up” RBDs (state 1; 3.4 Å resolution), each of which was complexed with a Fab, and a symmetric trimer with three RBDs in the same “up” conformation (state 2; 3.7 Å resolution), again with each RBD complexed with a Fab (Figure 5A). A subset of complexes in the cryo-EM structure of the S230 mAb bound to SARS-CoV S trimer were also found with three “up” RBDs bound to three Fabs (Walls et al., 2019), although in that structure, as in the C105-S structure, the majority of complexes had their RBDs in a two “up,” one “down” configuration.

The C105-RBD interfaces were similar across the five examples in the state 1 and state 2 complex structures (Figure S5), thus we describe the interface for one of the Fab-RBD complexes in the state 1 complex in which the resolution at the interface was improved by performing a focused refinement (Punjani et al., 2017) on the C105 Fab-RBD portion of the complex (Figure S5). The C105 Fab uses its three heavy chain complementarity determining regions (CDRH1, CDRH2, and CDRH3) and two of its light chain CDRs (CDRL1 and CDRL3) to rest against the receptor-binding ridge of the RBD (Figure 5B,C). The majority of the antibody contacts are made by CDRH1, CDRH2, and CDRL1, with CDRH3 and CDRL3 playing minor roles. The C105 epitope overlaps with the COV21 epitope defined by nsEMPEM, which also rests against the receptor-binding ridge in the RBD, although the Fab(s) in the COV21 reconstruction are predicted to adopt a different angle of approach (Figure 5D). Interestingly, the C105-RBD interaction closely resembles the RBD interaction of another COVID-19 donor-derived neutralizing mAb, B38 (Figure 5E), as reported in a recent Fab-RBD crystal structure (Wu et al., 2020c). The heavy chains of both B38 and C105 are derived from the *VH3-53* gene segment, whereas the light chain gene segments differ: *KV1-9* for B38 (Wu et al., 2020c) and *LV2-8* for C105 (Robbiani et al., 2020). Accordingly, the CDRH1 and CDRH2 loops of both neutralizing antibodies share similar conformations and contribute more to the antibody-RBD interface than their CDRH3 loops (Figure 5B,E).

The common epitope of C105 and B38 overlaps with the binding site for ACE2 (Figure 5F), rationalizing their potent neutralizing activities (Robbiani et al., 2020; Wu et al., 2020c). Given that COV21 was one of the more potent neutralizing plasmas of the 149 that were collected (Robbiani et al., 2020), the overlap in the C105/B38 neutralizing epitope with the nsEMPEM-defined predominant COV21 epitope suggests that recognition of the COV21 epitope by S230-like antibodies would also be neutralizing.

### CDRH3 length is a characteristic of the recurring *VH3-53*/*VH3-66* class of anti-SARS-CoV-2 RBD neutralizing antibodies

The shared binding mode of B38 and C105, both *VH3-53*-derived mAbs, defines a recurrent class of anti-SARS-CoV-2 mAbs. Among a large set (*n*=534) of cloned anti-SARS-CoV-2 mAbs against the RBD, those derived from *VH3-53* and *VH3-66* were over-represented (Robbiani et al., 2020). Other studies have also reported anti-SARS-CoV-2 mAbs derived from these genes (Brouwer et al., 2020; Cao et al., 2020; Chi et al., 2020; Ju et al., 2020; Rogers et al., 2020; Seydoux et al., 2020; Wu et al., 2020c; Zost et al., 2020). These VH gene segments encode V regions that differ in only one amino acid position, which is not in a CDR. Thus, in terms of V-gene-determined mAb classes, they are functionally equivalent. When grouping *VH3-53* and *VH3-66*, over-representation of *VH3-53*/*VH3-66*-derived mAbs is significant (*p* = 0.035). A notable characteristic of the *VH3-53*/*VH3-66*-derived subset (75 mAbs) within the 534 anti-RBD mAbs (Robbiani et al., 2020) was a bias towards shorter CDRH3s (as defined by IMGT) (Lefranc et al., 2015): 75% had lengths between 9 and 12 residues, which is significantly different from the human repertoire and from the entire set of 534 anti-SARS-CoV-2 RBD mAbs (two sample Kolmogorov-Smirnov test, *p* < 0.001) (Figure S7A).

Superposition of VH domains from unrelated antibodies with longer CDRH3s suggests that RBD residues 456-457 and/or 484-493 present a steric barrier limiting the CDRH3 loop lengths that are compatible with this binding orientation (Figure S7B). A recent report identified a clonally-unrelated group of *VH3-53*/*VH3-66* anti-SARS-CoV-2 mAbs based on CDRH3 sequence similarity to the anti-SARS-CoV mAb m396 (derived from *VH1-69*) (Cao et al., 2020). The structure of a SARS-CoV RDB complex with m396 (PDB code 2DD8) (Prabakaran et al., 2006) shows that mAb m396 does not share the B38/C105 binding mode. We suggest that the key feature of the *VH3-53*/*VH3-66* mAbs identified based on CDRH3 sequence similarity to m396 (Cao et al., 2020) is their CDRH3 length (11 residues), and that these mAbs will share the B38/C105 binding mode, not the m396 binding mode.

### Identified S mutations are unlikely to affect epitopes revealed by nsEMPEM and single-particle cryo-EM

A recent report suggested that a mutation in the S protein (D614G) increases transmissibility of SARS-CoV-2 (Korber et al., 2020), and it has been speculated that this substitution, or others found in different S protein sequences, could affect antibody recognition. In cryo-EM structures of the prefusion S trimer (Walls et al., 2020; Wrapp et al., 2020) and in our C105-S complex (Figure 5), S protein residue D614 is located in S1^D^, where it makes contact with an adjacent protomer. To address whether the D614G mutation could affect binding of antibodies in COV21 and COV57 plasma samples, we marked the location of the D614 residues and other residues that were reported to mutate (Korber et al., 2020) on the COV21 and COV57 nsEMPEM reconstructions (Figure 6A,B) and on the C105-S cryo-EM structure (Figure 6C). The RBD-binding COV21 Fab(s) and the C105 Fab are distant from residue D614 (Figure 6A,C). Therefore, if the COV21 reconstruction reflects the predominant epitope in the COV21 plasma, it is unlikely that antibodies elicited in the COV21 individual would be sensitive to the D614G substitution. Indeed, in the absence of large conformational changes, all anti-RBD antibodies, including C105 (Robbiani et al., 2020) and B38 (Wu et al., 2020c), would be unaffected by this substitution. Mutations in the SARS-CoV-2 RBD identified by genome sequencing also include N439K, V483A, and V367F (Tables S3,S4), but the affected residues are not within the epitopes of the COV21 Fab(s) (Figure 6A,B) or the C105 Fab (Figure 6C), and residue 483 is disordered in unliganded S protein structures (Walls et al., 2020; Wrapp et al., 2020). The predominant epitope in the COV57 plasma is closer to S protein D614, but this residue appears to be outside of the binding interface. In addition, mutations in the S1^A^ domain identified by genome sequencing (Tables S3,S4) reside outside of the COV57 Fab(s) epitope (Figure 6A,B). Thus in the absence of a major conformational change induced by mutation, the observed substitutions, particularly the D614G mutation, are unlikely to affect antibodies elicited in the COV21 or COV57 individuals or in RBD-binding antibodies such as C105.

**Figure 6.**
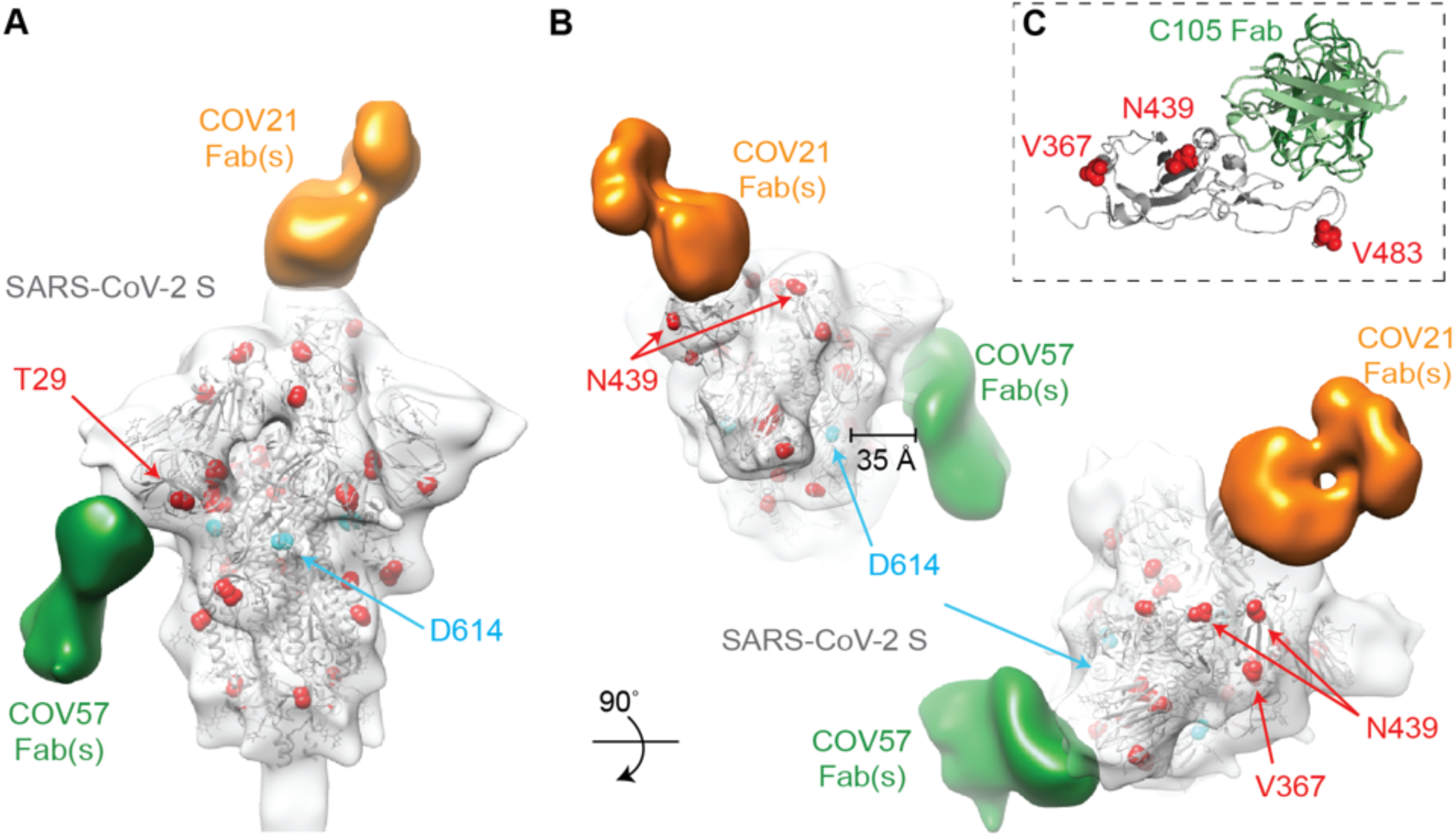
Identified S mutations are unlikely to affect epitopes revealed by nsEMPEM and single-particle cryo-EM. (A,B) Refined 3D model of SARS-CoV-2 S trimer alone was fitted with a reference structure (PDB 6VYB; gray cartoon) with locations of mutations observed in circulating SARS-CoV-2 isolates (Table S3) highlighted (red spheres). Residues affected by mutations that are disordered in the SARS-CoV-2 S structure (V483A) or in regions that are not included in the S ectodomain (signal sequence or cytoplasmic tail) are not shown. Densities corresponding to Fabs were separated, colored, and displayed on the same 3D volume. (C) C105-RBD interaction from the cryo-EM structure of the C105-S complex (Figure 5) showing locations of RBD mutations. V483 is ordered in this structure. See also Tables S3 and S4.

### S protein epitopes offer different possibilities for avidity effects during IgG and receptor binding

IgGs contain two identical antigen-binding Fabs, thus offering the opportunity to bind pathogens with regularly-spaced antigen sites using avidity effects, either through inter-spike crosslinking (binding the same epitope on adjacent spikes) and/or intra-spike crosslinking (binding the same epitope on identical subunits of a single multimeric spike) (Klein and Bjorkman, 2010).

To address whether inter-spike crosslinking by anti-SARS-CoV-2 IgGs could occur, we modeled adjacent S proteins on a virion membrane assuming a minimum inter-spike separation distance of ∼15 nm, as observed from cryo-electron tomography analyses of SARS-CoV and other coronaviruses (Neuman et al., 2011). By including a bound Fab on each S trimer in the position of the COV21 Fab(s) from the nsEMPEM reconstruction, we addressed whether the Fabs from a single IgG could bind to two adjacent S trimers. The modeling predicts that inter-spike crosslinking could occur for the COV21 epitope (Figure 7A). By contrast, the downward-pointing Fab(s) in the COV57-S nsEMPEM reconstruction appear unlikely to participate in inter-spike crosslinking by an IgG due to the Fab orientations being unable to accommodate an Fc in a position that could join two Fabs (Figure 7A). These predictions are consistent with ELISA results demonstrating diminished binding for COV21 Fabs compared with their IgG counterparts, but less pronounced differences for the Fab versus IgG comparison for COV57 (Figure 2).

**Figure 7.**
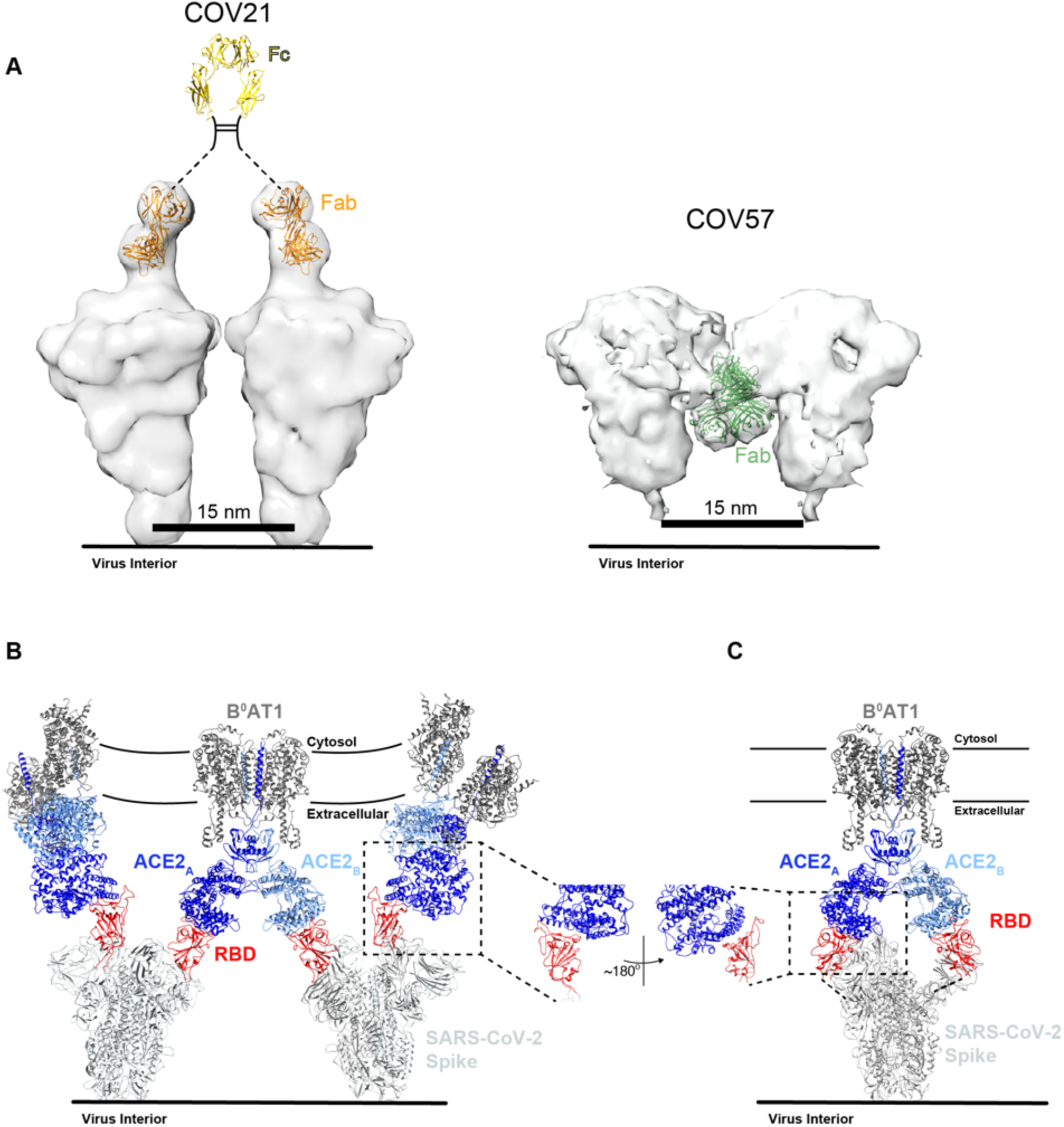
S protein epitopes offer different possibilities for avidity effects during IgG and receptor binding. (A) Left: Model of two adjacent S trimers separated by ∼15 nm, as seen on coronaviruses by cryo-electron tomography (Neuman et al., 2011), demonstrating that the orientation of COV21 Fab(s) on S could accommodate inter-spike crosslinking by a single IgG. The Fc portion of the IgG (PDB code 1IGT) was modeled assuming flexibility between the Fabs and the Fc (Sandin et al., 2004) and with the hinge region indicated by a dotted line since it is disordered in crystal structures of intact IgGs (Harris et al., 1992; Harris et al., 1998; Saphire et al., 2001). Right: Example of a model of two adjacent S trimers with bound Fab(s) in the orientation observed in the COV57 Fab(s)-S reconstruction demonstrating that inter-spike crosslinking is unlikely due to the “downward” orientation of the Fab(s), which does not permit linking by an Fc region, and predicted steric clashes between adjacent Fabs. Inter-spike crosslinking is also not possible for other orientations of two adjacent COV57 Fab(s)-S complexes (not shown). (B) Model of S trimers with two RBDs in an “up” position based on a cryo-EM structure of SARS-CoV S trimer (Kirchdoerfer et al., 2018) (PDB code 6CRX) interacting with full-length ACE2 receptors from the cryo-EM structure of soluble SARS-CoV-2 RBDs bound to the dimeric membrane form of ACE2 (Yan et al., 2020) (PDB code 6M17). Inter-spike crosslinking is possible if ACE2 dimers cluster in the membrane. (C) Model of intra-spike crosslinking between dimeric ACE2 and an S protein trimer with two RBDs in an “up” position. The RBDs were rotated by ∼180° about their long axes to allow binding of the ACE2 ectodomains. Rotation of the RBD is a possibility since its position is flexible with respect to the remaining part of the S trimer (Walls et al., 2019). In this model, RBDs from a single S trimer could bind the ACE2 dimer in the same configuration as seen in the B°AT1–ACE2–SARS-CoV-2 RBD structure (PDB code 6M17).

We also used modeling to predict whether avidity effects could influence the interaction between ACE2, an integral membrane protein that dimerizes on the target cell surface (Yan et al., 2020), with viral S trimers. Starting with a cryo-EM structure of dimeric full-length ACE2 associated with the integral membrane protein B°AT1 bound to monomeric SARS-CoV-2 RBDs, we modeled bound S trimers from a cryo-EM structure of SARS-CoV S trimers with two RBDs in an “up” position (Kirchdoerfer et al., 2018). Assuming that there are adjacent ACE2-B°AT1 complexes in the host cell membrane, the modeling predicts that inter-spike crosslinking is possible (Figure 7B). Assuming rotation of the RBDs in a two “up”/one “down” S trimer, intra-spike cross-linking could also occur (Figure 7C). If S trimers can indeed crosslink adjacent ACE2 receptors or bind as a single trimer to both ACE2 subunits in an ACE2 dimer, they could take advantage of avidity effects to bind more tightly than predicted from affinity measurements involving the interactions of monomeric ACE2 ectodomains to monomeric coronavirus RBDs (Shang et al., 2020; Walls et al., 2020; Wrapp et al., 2020).

The possibility of avidity effects during the interactions of SARS-CoV-2 S with ACE2 dimers has implications for interpretation of pseudovirus assays to measure coronavirus infectivity in the presence and absence of potential inhibitors such as antibodies. In vitro neutralization assays for SARS-CoV-2 include pseudoviruses based on HIV lentiviral particles (Chen et al., 2020; Crawford et al., 2020; Ou et al., 2020; Robbiani et al., 2020; Wu et al., 2020a), murine leukemia virus retroviral particles (Pinto et al., 2020; Quinlan et al., 2020), and vesicular stomatitis virus (Hoffmann et al., 2020; Nie et al., 2020; Xiong et al., 2020). Each of these pseudovirus types could potentially incorporate different numbers of spikes, in which case the overall spike density would alter sensitivity to antibody avidity. In any case, the effects of avidity on IgG binding to a tethered antigen are a complicated mixture of intrinsic Fab-antigen affinity, kinetics, input concentration, and incubation time (Klein and Bjorkman, 2010; Wu et al., 2005), thus neutralization potencies of some, but not all, IgGs could be affected in *in vitro* neutralization assays. In addition, when considering therapeutic applications of convalescent plasma or purified antibodies, avidity effects would be difficult to predict given uncertainties about antibody concentrations, viral titers, and potentially different S trimer spacings and densities on infectious virions.

## Conclusions

Our results provide a glimpse into diverse antibody responses in neutralizing plasmas from donors who recovered from COVID-19. We characterized polyclonal plasma IgGs that exhibited different degrees of cross-reactive binding between S proteins from SARS-CoV-2, SARS-CoV, and MERS-CoV and showed that the plasma IgGs also included non-cross-reactive antibodies against common cold virus RBDs. By mapping SARS-CoV-2 S epitopes targeted by convalescent plasma IgGs, we not only observed the expected targeting of the S protein RBD, but also discovered an epitope outside of the RBD, which may represent an alternative binding site for neutralizing antibodies. The RBD-binding Fab(s) from COV21 plasma resembled binding of S230, a *VH3-30* mAb isolated from a SARS-CoV patient that blocks ACE2 receptor binding (Rockx et al., 2008). We found another type of ACE2 receptor-blocking anti-SARS-CoV-2 antibody in our analysis of a neutralizing mAb derived from the COV107 individual. In a 3.4 cryo-EM structure SARS-CoV-2 S protein bound to this mAb, C105, we observed an epitope on the RBD that overlapped with the binding site for COV21 Fab(s) and closely resembled the binding of another mAb, B38 (Wu et al., 2020c). Like C105, B38 is also derived from the *VH3-53* VH gene segment. Our structural studies support the hypothesis that recurrent classes of anti-SARS-CoV-2 neutralizing antibodies derived from the *VH3-53/VH3-66* and *VH3-30* gene segments use the distinct RBD-binding modes of the B38/C105 and S230 mAbs, respectively, providing valuable information for evaluating antibodies raised by infection or vaccination by sequences alone. Finally, the RBD and S1^A^ epitopes we mapped by nsEMPEM and single-particle cryo-EM are unlikely to be affected by common mutations in different SARS-CoV-2 isolates, offering hope that antibody therapeutics and/or a vaccine might be effective in combatting the current pandemic.

## Acknowledgements

We thank all study participants who devoted time to our research, Drs. Barry Coller and Sarah Schlesinger, and the Rockefeller University Hospital Clinical Research Support Office and nursing staff. We thank the Global Initiative on Sharing Avian Influenza Data (GISAID) and the originating and submitting laboratories for sharing the SARS-CoV-2 genome sequences; see Table S4 for a list of sequence contributors. We thank members of the Bjorkman, Nussenzweig, and Bienasz laboratories for helpful discussions, Drs. John Pak (Chan Zuckerberg Biohub) and Florian Krammer (Icahn School of Medicine at Mount Sinai) for CoV expression plasmids, and Drs. Songye Chen and Andrey Malyutin (Caltech) for maintaining electron microscopes. Electron microscopy was performed in the Caltech Beckman Institute Resource Center for Transmission Electron Microscopy. We thank the Gordon and Betty Moore and Beckman Foundations for gifts to Caltech to support the Molecular Observatory (Dr. Jens Kaiser, director), and Drs. Silvia Russi, Aina Cohen, and Clyde Smith and the beamline staff at SSRL for data collection assistance. This work was supported by NIH grant P01-AI138398-S1 (P.J.B. and M.C.N.), the Caltech Merkin Institute for Translational Research (P.J.B.), NIH grant P50 8 P50 AI150464-13 (P.J.B.), 2U19AI111825 (M.C.N.), and George Mason University Fast Grants (P.J.B., D.F.R.). C.O.B was supported by the Hanna Gray Fellowship Program from the Howard Hughes Medical Institute and the Postdoctoral Enrichment Program from the Burroughs Wellcome Fund. C.G. is a fellow of the Robert S. Wennett and Mario Cader-Frech Foundation and was supported in part by grant # UL1 TR001866 from the National Center for Advancing Translational Sciences (NCATS, National Institutes of Health Clinical and Translational Science Award (CTSA) program) and by the Shapiro-Silverberg Fund for the Advancement of Translational Research). P.D.B. and M.C.N. are Howard Hughes Medical Institute Investigators.

## Author contributions

C.O.B., A.P.W., D.F.R., M.C.N., and P.J.B. conceived the study and analyzed data. C.O.B. performed protein characterization, nsEMPEM and cryo-EM, and interpreted structures. N.G.S. and C.O.B. performed and analyzed crystallographic structures. C.O.B. and K.H.T. prepared plasma IgGs and Fabs. M.A.G.H., K.H.T., N.G.S., and C.O.B. performed and analyzed ELISAs. P.R.H. and N.K. expressed and purified proteins. A.P.W. and H.B.G. performed computational modeling. A.P.W. analyzed antibody sequences. F.M., J.C.C.L., Y.W., F.S., T.H., and P.D.B. developed and performed in vitro neutralization assays. C.G., S.F., A.H., K.G.M., M.C., and D.F.R. collected and processed human plasma samples. C.O.B., A.P.W., M.C.N., and P.J.B. wrote the paper with contributions from other authors.

## Declaration of Interests

The authors declare no competing interests.

## CONTACT FOR REAGENT AND RESOURCE SHARING

Further information and requests for reagents should be directed to and will be fulfilled by the Lead Contact, Pamela Bjorkman for reagents not related to human samples or pseudovirus. For information or requests concerning human samples, please contact Davide Robbiani and Michel Nussenzweig; for pseudoviruses and neutralization assays, please contact Paul Bieniasz. Sharing of reagents with academic researchers may require UBMTA agreements.

## EXPERIMENTAL MODEL AND SUBJECT DETAILS

### Human subjects

Samples of peripheral blood were obtained upon written consent from community participants under protocols approved by the Institutional Review Board of the Rockefeller University (DRO-1006). Details on the demographics of the cohort are provided in (Robbiani et al., 2020).

### Cell lines

HEK293T cells for pseudovirus production and HEK293T_ACE2_ cells for pseudovirus neutralization experiments were cultured at 37°C and 5% CO_2_ in Dulbecco’s modified Eagle’s medium (DMEM, Gibco) supplemented with 10% heat-inactivated fetal bovine serum (FBS, Sigma-Aldrich) and 5 µg/ml Gentamicin (Sigma-Aldrich). The medium for the 293T_Ace2_ cells additionally contained 5µg/ml Blasticidin (Gibco). For constitutive expression of ACE2 in 293T cells, a cDNA encoding ACE2, carrying two inactivating mutations in the catalytic site (H374N & H378N), was inserted into CSIB 3’ to the SFFV promoter (Kane et al., 2016). 293T_ACE2_ cells were generated by transduction with CSIB based virus followed by selection with 5 μg/ml Blasticidin (Gibco). Expi293F cells (Gibco) for protein expression were maintained at 37°C and 8% CO_2_ in Expi293 Expression medium (Gibco), transfected using Expi293 Expression System Kit (Gibco) and maintained under shaking at 130 rpm. The gender of the HEK293T, HEK293T_ACE2_ and Expi293F cell lines is female. Cell lines were not specifically authenticated.

### Bacteria

*E. coli* DH5*α* (Zymo Research) for propagation of expression plasmids were cultured at 37°C in LB broth (Sigma-Aldrich) with shaking at 250 rpm.

### Viruses

To generate pseudotyped viral stocks, HEK293T cells were transfected with pNL4-3ΔEnv-nanoluc and pSARS-CoV2-S_trunc_ (Robbiani et al., 2020) using polyethyleneimine, leading to production of HIV-1-based virions carrying the SARS-CoV-2 S protein at the surface. Eight hours after transfection, cells were washed twice with PBS and fresh media was added. Supernatants containing virions were harvested 48 hours post transfection, filtered and stored at −80°C. Infectivity of virions was determined by titration on 293T_ACE2_ cells.

## METHOD DETAILS

### Collection of human samples

Convalescent and healthy donor plasma samples were collected and processed as described (Robbiani et al., 2020). The convalescent plasma samples used for nsEMPEM were from residents in the State of New York: COV21 (a 54-year-old male Hispanic, collection 27 days after symptom onset), COV57 (a 66-year-old male Caucasian, collection 21 days after symptom onset), and COV107 (a 53-year-old female Caucasian, collection 29 days after symptom onset). An analysis of SARS-CoV-2 genomes in the GISAID database with sample collection dates in March 2020 (contemporaneous with the infections of individuals COV21, COV57, and COV107) was performed to identify any viral spike mutations likely to have been present. For SARS-CoV-2 genomes of New York State residents from March 2020, 468 of 475 contained the D614G mutation. Thus, based on state-level mutant frequencies, these individuals were likely to have been infected with D614G-containing viruses. All other spike mutations in these genomes had a frequency below 2%. All participants provided written informed consent before sample collection at the Rockefeller University Hospital and the study was conducted in accordance with Good Clinical Practice. Anti-coagulated plasma was heat-inactivated (56°C for 1 hour) prior to shipment to Caltech and stored at 4°C thereafter (Robbiani et al., 2020).

### Phylogenetic trees

Sequence alignments of S proteins and RBD/S1^B^ domains were made with Clustal Omega (Sievers et al., 2011). Phylogenetic trees were calculated from these amino acid alignments using PhyML 3.0 (Guindon et al., 2010) and visualized with PRESTO (http://www.atgc-montpellier.fr/presto).

### Cloning and expression of recombinant CoV proteins

Codon-optimized nucleotide sequences encoding the SARS-CoV-2 S ectodomain (residues 16-1206 of an early SARS-CoV-2 sequence isolate; GenBank MN985325.1, which has an Asp at position 614, so does not include the D614G mutation described as possibly more transmissible in (Korber et al., 2020)), SARS-CoV S (residues 12-1193; GenBank AAP13441.1), MERS-CoV S (residues 19-1294; GenBank JX869059.2), HCoV-OC43 (residues 15-1263; GenBank AAT84362.1), HCoV-NL63 (residues 16-1291; GenBank AAS58177.1), and HCoV-229E (residues 17-1113; GenBank AAK32191.1) were synthesized and subcloned into the mammalian expression pTwist-CMV BetaGlobin vector by Twist Bioscience Technologies. The S proteins were modified as previously described (Li et al., 2019; Tortorici et al., 2019; Walls et al., 2020). Briefly, the S ectodomain constructs included an N-terminal mu-phosphatase signal peptide, 2P stabilizing mutations (Pallesen et al., 2017) and a C-terminal extension (GSG-RENLYFQG (TEV protease site), GGGSG-YIPEAPRDGQAYVRKDGEWVLLSTFL (foldon trimerization motif), G-HHHHHHHH (octa-histidine tag), and GLNDIFEAQKIEWHE (AviTag)). For SARS-CoV-2, MERS-CoV, HCoV-NL63 and HCoV-OC43 mutations to remove the S1/S2 furin cleavage site were introduced.

Codon-optimized nucleotide sequences encoding the receptor binding domain (RBD) for SARS-CoV-2 (residues 331-524) SARS-CoV S (residues 318-510), MERS-CoV S (residues 367-588), HCoV-NL63 (residues 481-614), and corresponding S1^B^ domains for HCoV-OC43 (residues 324-632), and HCoV-229E (residues 286-434) were synthesized and subcloned into the mammalian expression pTwist-CMV BetaGlobin vector by Twist Bioscience Technologies. The RBD/S1^B^ constructs included an N-terminal human IL-2 signal peptide and dual C-terminal tags (G-HHHHHHHH (octa-histidine), and GLNDIFEAQKIEWHE (AviTag)).

The S protein and RBD/S1^B^ constructs were expressed by transient transfection of Expi293F cells (Gibco) and purified from clarified transfected cell supernatants four days post-transfection using HisTrap FF and HisTrap HP columns (GE Healthcare Life Sciences). After concentration, affinity-purified proteins were further purified by size-exclusion chromatography (SEC) using a HiLoad 16/600 Superdex 200 pg column (GE Healthcare Life Sciences) in 1x TBS (20 mM Tris-HCl pH 8.0, 150 mM NaCl, 0.02% NaN_3_). Peak fractions were analyzed by SDS-PAGE, and fractions corresponding to S trimers or monomeric RBD/S1^B^ proteins were pooled and stored at 4°C.

### SEC-MALS

Purified CoV-S trimers were concentrated to 1 mg/mL and loaded onto a Superose 6 Increase 10/300 GL column (GE Healthcare Life Sciences) and passed through a Wyatt DAWN coupled to a Wyatt UT-rEX differential refractive index detector (Wyatt Technology). Data were analyzed using Astra 6 software (Wyatt Technology) to determine glycoprotein molecular weights.

### Purification of plasma IgGs and Fabs

IgGs were purified from plasma samples using 5-mL HiTrap MabSelect SuRe columns (GE Healthcare Life Sciences). Heat-inactivated plasma was diluted 10-fold with cold PBS, and samples were applied to prepacked columns at 1 mL/min. Bound IgGs were washed with 10 column volumes (CV) PBS and eluted with 5 CV 0.1M glycine, 100 mM NaCl, pH 3.0 directly into 10% v/v 1M Tris-HCl, pH 8. To produce polyclonal Fab fragments, IgGs were buffer exchanged into PBS by centrifugation with 30 kDa MWCO membrane centrifugal filter units (Millipore). Fabs were generated by papain digestion using crystallized papain (Sigma-Aldrich) in 50 mM sodium phosphate, 2 mM EDTA, 10 mM L-cysteine, pH 7.4 for 30-60 min at 37°C at a 1:100 enzyme:IgG ratio. To remove undigested IgGs and Fc fragments, digested products were applied to a 1-mL HiTrap MabSelect SuRe column (GE Healthcare Life Sciences) and the flow-through containing cleaved Fabs was collected. Fabs were further purified by SEC using a Superdex 200 Increase 10/300 column (GE Healthcare Life Sciences) in TBS, before concentrating and storage at 4°C.

To evaluate binding of purified polyclonal IgGs or Fabs to CoV proteins, purified S1^B^/RBD or S proteins were biotinylated using the Biotin-Protein Ligase-BIRA kit according to manufacturer’s instructions (Avidity). Biotinylated-CoV proteins were captured on streptavidin coated 96-well plates (Thermo Scientific) by incubating with 100 µL of 2 µg/mL protein solution in TBS overnight at 4°C. Plates were washed 3x in washing buffer (1x TBST: 20 mM Tris, 150 mM NaCl, 0.05% Tween20, pH 8.0) and blocked with 200 µL blocking buffer (TBST-MS: 1x TBST + 1% w/v milk, 1% v/v goat serum (Gibco) for 1 h at RT. Immediately after blocking, polyclonal IgGs or Fabs were assayed for binding at a 50µg/mL starting concentration and seven 4-fold serial dilutions in blocking buffer. After 2 h incubation at RT, plates were washed 5 times with washing buffer and incubated with goat-anti-human IgG or goat-anti-human IgG(H+L) secondary antibody conjugated to horseradish peroxidase (HRP) (SouthernBiotech) in blocking buffer at a 1:4000 or 1:2000 dilution, respectively. Plates were washed 5 times with washing buffer and developed by addition of 100µL 1-Step™ Ultra TMB-ELISA Substrate Solution (Thermo Scientific) for 3 min. The developing reaction was quenched by addition of 100µl 1N HCl and absorbance was measured at 450nm using Gen5 software on a Synergy Neo2 Reader (BioTek).

### RBD Adsorption ELISAs

Plasmids for SARS-CoV-2 RBD 6xHisTag constructs (residues 319-541, GenBank: MN908947.3) were a gift from the lab of Florian Krammer (Mount Sinai). SARS-CoV-2 RBD 6xHisTag constructs were expressed by transient transfection of Expi293F cells (Gibco) and purified using HisTrap FF and HisTrap HP columns (GE Healthcare Life Sciences), followed by SEC using a HiLoad 16/600 Superdex 200 pg column (GE Healthcare Life Sciences) against 1x TBS (20 mM Tris-HCl pH 8.0, 150 mM NaCl, 0.02% NaN_3_). Purified protein was concentrated and buffer exchanged into 100 mM Sodium Bicarbonate pH 8.3, 500 mM NaCl using a gravity-flow chromatography with a PD-10 desalting column (GE Healthcare Life Sciences). Buffer-exchanged RBD was concentrated to 5 mL and covalently coupled to a 5 mL HiTrap NHS-activated Sepharose column (GE Healthcare Life Sciences) according to the manufacturer’s protocol. Control resin was made using the same procedure to covalently couple 2G12, an HIV-1 mAb, as described (Scharf et al., 2015).

For RBD-absorption ELISA experiments to evaluate binding of purified polyclonal IgGs to CoV S1^B^/RBD proteins after absorption of SARS-CoV-2 RBD-specific antibodies, biotinylated-S1^B^/RBD proteins were captured on high-capacity streptavidin coated 96-well plates (Thermo Scientific) by incubating with 100 µL of 2 µg/mL protein solution in TBS overnight at 4°C. Plates were washed 3x in washing buffer and blocked as described above. For absorption of RBD-specific antibodies, 100 µL of SARS-CoV-2 RBD-coupled resin or 2G12 control resin was incubated with 100 µL of purified IgGs diluted to ∼1 mg/mL for 1 h at RT with agitation. After incubation, SARS-CoV-2 RBD-coupled resin was gently centrifuged at 250 x *g* for 2 min, and non-absorbed IgGs were removed by careful pipetting of the aqueous layer above the pelleted RBD-coupled resin. Unadsorbed and absorbed IgG samples were assayed at a 50 µg/mL starting concentration and seven 4-fold serial dilutions as described above.

### Pseudotyped virus neutralization assays

Pseudoviruses based on HIV lentiviral particles were prepared as described (Robbiani et al., 2020). Four-fold serially diluted purified polyclonal IgGs and Fabs from COVID-19 plasmas were incubated with SARS-CoV-2 pseudotyped virus for 1 hour at 37°C. After incubation with 293T_ACE2_ cells for 48 hours at 37°C, cells were washed twice with PBS, lysed with Luciferase Cell Culture Lysis 5x reagent (Promega), and NanoLuc Luciferase activity in lysates was measured using the Nano-Glo Luciferase Assay System (Promega). Relative luminescence units (RLUs) were normalized to values derived from cells infected with pseudotyped virus in the absence of purified plasma IgGs or Fabs. Half-maximal inhibitory concentrations (IC_50_ values) for purified plasma IgGs and Fabs were determined as molar concentrations (to account for the IgG versus Fab difference in molecular weight) using 4-parameter nonlinear regression (Prism, GraphPad).

### Negative-stain electron microscopy (nsEM)

Purified CoV-S trimers were adsorbed to freshly-glow discharged PureC 300 mesh carbon-coated copper grids (EMD Sciences) for 1 min followed by 2% uranyl formate staining. Micrographs were recorded using Digital Micrograph software on a 120kV FEI Tecnai T12 equipped with a Gatan Ultrascan 2k X 2k CCD at a 52,000x nominal magnification.

### nsEMPEM

Methods were adapted from (Bianchi et al., 2018). To form polyclonal Fab-S complexes, 30 µg of SARS-CoV-2 S trimers were incubated overnight at RT with 30-50 mg/mL Fabs in 100 µL total volume (corresponding to ∼1000x the EC_50_ values for ELISAs using purified polyclonal Fabs), and the complexes were purified by SEC on a Superose 6 increase 10/300 GL column (GE Healthcare Life Sciences). Fractions containing complexes were pooled and concentrated to 50 µg/mL and passed through a 0.1 µm filter before deposition on 300 mesh carbon-coated copper grids (source?) and stained with 1% (w/v) uranyl formate (source?). Grids were imaged at 300 keV using a Titan Krios transmission electron microscope (Thermo Fisher) operating at RT, equipped with a K3 direct electron detector (Gatan) using SerialEM 3.7? (Mastronarde, 2005). Images were processed in cryoSPARC v 2.14, and a reference-free particle stack was generated using a Gaussian blob picker (Punjani et al., 2017). Particles corresponding to S-Fab complexes were identified by extensive 2D classification to identify class averages that displayed structural elements interpreted as Fab density and also represented different views. Extracted particles were used to generate ab initio models in cryoSPARC that were further processed by 3D classification to separate out complexes and S trimer structures alone. Figures were prepared using UCSF Chimera (Goddard et al., 2007; Pettersen et al., 2004).

### X-ray crystallography

The Fab from the C105 monoclonal IgG was expressed, purified, and stored as described (Scharf et al., 2015; Schoofs et al., 2019). Crystallization trials were performed at room temperature using the sitting drop vapor diffusion method by mixing equal volumes of C105 Fab and reservoir using a TTP LabTech Mosquito robot and commercially-available screens (Hampton Research). After optimization of initial hits, crystals were obtained in 0.15 M lithium sulfate, 0.1 M citric acid pH 3.5, 18% v/v PEG 6000 at 20 °C. Crystals were transferred stepwise to 20% glycerol cryoprotectant before being cryopreserved in liquid nitrogen.

X-ray diffraction data were collected for C105 Fab at the Stanford Synchroton Radiation Lightsource (SSRL) beamline 12-1 on a Pilatus 6M pixel detector (Dectris). Data from a single crystal were indexed and integrated in XDS (Kabsch, 2010) and merged using AIMLESS in *CCP4* (Winn et al., 2011) (Table S1). The structure of C105 Fab was determined by molecular replacement in PHASER (McCoy et al., 2007) using the B38 Fab coordinates from PDB code 7BZ5 after removal of CDR loops as a search model. The C105 Fab coordinates were refined using Phenix (Adams et al., 2010) and cycles of manual building in Coot (Emsley et al., 2010) (Table S1).

### Cryo-EM Sample Preparation

Purified C105 Fab was incubated with SARS-CoV-2 S trimer at a 2:1 molar ratio per protomer on ice for 30 minutes prior to purification by SEC on a Superose 6 Increase 10/300 GL column (GE Healthcare Life Sciences). Fab–S complexes were concentrated to 1.6 mg/ml in Tris-buffered saline (TBS). Immediately before deposition onto a 300 mesh, 1.2/1.3 UltraAuFoil grid (Electron Microscopy Sciences) that had been freshly glow-discharged for 1 min at 20 mA using a PELCO easiGLOW (Ted Pella), 3 μL of complex was mixed with 0.5 µL of a 0.5% w/v octyl-maltoside solution (Anatrace). Samples were vitrified in 100% liquid ethane using a Mark IV Vitrobot (Thermo Fisher) after blotting for 3 s with Whatman No. 1 filter paper at 22°C and 100% humidity.

### Cryo-EM Data Collection and Processing

For the C105-S trimer complex, micrographs were collected on a Titan Krios transmission electron microscope (Thermo Fisher) operating at 300 kV using SerialEM automated data collection software (Mastronarde, 2005). Movies were obtained on a Gatan K3 Summit direct electron detector operating in counting mode at a nominal magnification of 105,000x (super-resolution 0.418 Å/pixel) using a defocus range of −1 to −2.5 μm. Movies were collected with an 1.9 s exposure time with a rate of 22 e^-^/pix/s, which resulted in a total dose of ∼60 e-/Å^2^ over 40 frames. The 5,940 cryo-EM movies were patch motion corrected for beam-induced motion including dose-weighting within cryoSPARC v2.15 (Punjani et al., 2017) after binning super resolution movies by 2 (0.836 Å/pixel). The non-dose-weighted images were used to estimate CTF parameters using CTFFIND4 (Rohou and Grigorieff, 2015), and micrographs with power spectra that showed poor CTF fits or signs of crystalline ice were discarded, leaving 5,316 micrographs. Particles were picked in a reference-free manner using Gaussian blob picker in cryoSPARC (Punjani et al., 2017). An initial 565,939 particle stack was extracted, binned x2 (1.68 Å/pixel), and subjected to iterative rounds of reference-free 2D classification to identify class averages corresponding to intact S-trimer complexes with well-defined structural features. This routine resulted in a new particle stack of 71,289 particles, which were unbinned (0.836 Å/pixel) and re-extracted using a 352 box size. Two *ab initio* volumes were generated, with one class of 61,737 particles revealing an S-trimer complexed with two C105 Fabs.

Particles were further 3D classified (k=3) and heterogeneously refined to reveal two distinct states of the C105-S trimer complex. State 1 (37,615 particles) displaying 2 “up” RBD conformations bound by 2 C105 Fabs, and state 2 (14,119 particles) that displayed 3 “up” RBD conformations bound by 3 C105 Fabs. Particles from states 1 and 2 were separately refined using non-uniform 3D refinement imposing either C1 or C3 symmetry, respectively, to final resolutions of 3.6 Å (state 1; 37,615 particles) and 3.7 Å (state 2; 14,119 particles) according to the gold-standard FSC (Bell et al., 2016). Given that the RBD “up” conformations with C105 Fabs bound were similar in both states 1 and 2, improvements to the resolution at the RBD-C105 Fab interface were achieved by combining the entire particle stack (∼52k particles) for a focused, non-uniform 3D refinement. A soft mask was generated from the state 1 volume (5-pixel extension, 10-pixel soft cosine edge) for the S1 subunits and C105 Fab variable domains. These efforts resulted in a modest improvement in the RBD-C105 Fab interface (Figure S6D), and an overall resolution of 3.4 Å according to the gold-standard FSC.

### Cryo-EM Structure Modeling and Refinement

Initial coordinates were generated by docking individual chains from reference structures into cryo-EM density using UCSF Chimera (Goddard et al., 2007). The following coordinates were used: SARS-CoV-2 S trimer: PDB code 6VYB, “up” RBD conformations: PDB code 7BZ5, C105 Fab: this study. These initial models were then refined into cryo-EM maps using one round of rigid body refinement followed by real space refinement. Sequence-updated models were built manually in Coot (Emsley et al., 2010) and then refined using iterative rounds of refinement in Coot and Phenix (Adams et al., 2010). Glycans were modeled at possible N-linked glycosylation sites (PNGSs) in Coot using ‘blurred’ maps processed with a variety of B-factors (Terwilliger et al., 2018). Validation of model coordinates was performed using MolProbity (Chen et al., 2010) and is reported in Table S3.

### Structural Analyses

Structural figures were made using PyMOL (Version 1.8.2.1 Schrodinger, LLC) or UCSF Chimera (Goddard et al., 2007). Local resolution maps were calculated using cryoSPARC v 2.15 (Punjani et al., 2017).

## DATA AND SOFTWARE AVAILABILITY

Coordinates for the C105 Fab and the C105-S complex will be deposited in the PDB. Cryo-EM reconstructions of the C105-S complex will made available through the EMDB.

## Notes

### Competing Interest Statement

The authors have declared no competing interest.

