## Supplemental_Information for "Structures of human antibodies bound to SARS-CoV-2 spike reveal common epitopes and recurrent features of antibodies"

**Table S1. Crystallographic data collection and refinement statistics for C105 Fab structure (related to Figure 5)**.

**Table S2. Cryo-EM data collection and refinement statistics for C105-S complex structure (related to Figure 5)**.

**Table S3. S protein mutations found in different SARS-CoV-2 isolates (related to Figure 6)**. List of SARS-CoV-2 spike mutations with a frequency ≥ 0.1% in a set of 15335 isolates downloaded from the Global Initiative for Sharing All Influenza Data (GISAID) SARS-CoV-2 sequence database on 5/3/20 (Elbe and Buckland-Merrett, 2017; Shu and McCauley, 2017). The genomes were processed with the nextstrain augur pipeline (https://github.com/nextstrain/augur) (Hadfield et al., 2018), using MAFFT v7.464 (Katoh and Standley, 2013) for sequence alignment and FastTree (Price et al., 2010) to generate a phylogenetic tree. The resulting data were then analyzed with a custom Swift program.

**Table S4. List of GISAID accession IDs, virus names, originating labs, submitting labs, and authors (related to Figure 6).** We gratefully acknowledge all labs and authors for making SARS-CoV-2 genome sequence data available for this research.

**Table S1. Crystallographic data collection and refinement statistics for C105 Fab structure (related to Figure 5)**.

| | C105 Fab *(12-1, SSRL)* |
|---|---|
| **PDB ID** | **XXXX** |
| **Data collection**[a] | |
| Space group | I222 |
| Unit cell (Å) | 67.4, 120.1, 123.3 |
| α, β, γ (°) | 90, 90, 90 |
| Wavelength (Å) | 1.0 |
| Resolution (Å) | 38.9-1.80 (1.84-1.80) |
| Unique Reflections | 46,713 (2752) |
| Completeness (%) | 100 (99.8) |
| Redundancy | 6.8 (6.5) |
| $CC_{1/2}$ (%) | 98.8 (54.1) |
| <I/σI> | 5.7 (1.2) |
| Mosaicity (˚) | 0.19 |
| $R_{merge}$ (%) | 18.1 (157) |
| $R_{pim}$ (%) | 7.9 (70.5) |
| Wilson *B* -factor | 16.8 |
| **Refinement and Validation** | |
| Resolution (Å) | 38.9 - 1.80 |
| Number of atoms | |
| Protein | 3,132 |
| Ligand | 10 |
| Waters | 477 |
| $R_{work}/R_{free}$ (%) | 18.7/21.6 |
| R.m.s. deviations | |
| Bond lengths (Å) | 0.006 |
| Bond angles (˚) | 0.853 |
| MolProbity score | 1.29 |
| Clashscore (all atom) | 4.2 |
| Poor rotamers (%) | 0 |
| Ramachandran plot | |
| Favored (%) | 97.6 |
| Allowed (%) | 2.4 |
| Disallowed (%) | 0 |
| Average *B* -factor (Å) | 27.1 |

[a]Numbers in parentheses correspond to the highest resolution shell

**Table S2. Cryo-EM data collection and refinement statistics for C105-S complex structure (related to Figure 5).**

| | **C105 SARS-CoV-2 S 2P (state 1)** | **C105 SARS-CoV-2 S 2P (state 2)** |
|---|---|---|
| **PDB** | | |
| **EMD** | | |
| Microscope | Titan Krios | Titan Krios |
| Camera | Gatan K3 Summit | Gatan K3 Summit |
| Magnification | 105,000x | 105,000x |
| Voltage (kV) | 300 | 300 |
| Recording mode | counting | counting |
| Dose rate ($e^-$/pixel/s) | 22.1 | 22.1 |
| Electron dose ($e^-$/Å$^2$) | 60 | 60 |
| Defocus range (μm) | 1.0 - 2.5 | 1.0 - 2.5 |
| Pixel size (Å) | 0.418 (super resolution); 0.836 (binned) | 0.418 (super resolution); 0.836 (binned) |
| Micrographs collected | 5,940 | 5,940 |
| Micrographs used | 5,336 | 5,336 |
| Total extracted particles | 71,289 | 71,289 |
| Refined particles | 37,615 | 14,119 |
| Symmetry imposed | C1 | C3 |
| Nominal Resolution (Å) | | |
| FSC 0.5 (unmasked/masked) | 3.90/3.60 | 4.30/3.90 |
| FSC 0.143 (unmasked.masked) | 3.40/3.20 | 3.70/3.50 |
| Map sharpening $B$-factor | | |
| **Refinement and Validation** | | |
| Number of atoms | | |
| Protein | 25,973 | |
| Ligand | 711 | |
| MapCC (global/local) | 0.86/0.84 | |
| R.m.s. deviations | | |
| Bond lengths (Å) | 0.008 | |
| Bond angles (˚) | 0.812 | |
| MolProbity score | 2.17 | |
| Clashscore (all atom) | 13.6 | |
| Poor rotamers (%) | 0.04 | |
| Ramachandran plot | | |
| Favored (%) | 90.9 | |
| Allowed (%) | 9 | |
| Disallowed (%) | 0.1 | |

**Table S3. S protein mutations found in different SARS-CoV-2 isolates (related to Figure 6)**.

| Mutation | Count | Frequency(%) | Location |
|---|---|---|---|
| D614G | 9688 | 63.2 | S1 domain D |
| P1263L | 115 | 0.7 | S2 cytoplasmic tail |
| L5F | 91 | 0.6 | signal sequence |
| D936Y | 88 | 0.6 | S2 HR1 |
| L54F | 58 | 0.4 | S1 domain A |
| G1124V | 56 | 0.4 | S2 |
| N439K | 38 | 0.2 | S1 domain B (RBD) |
| H49Y | 35 | 0.2 | S1 domain A |
| L18F | 31 | 0.2 | S1 domain A |
| L8V | 30 | 0.2 | signal sequence |
| A831V | 29 | 0.2 | S2 |
| D839Y | 28 | 0.2 | S2 |
| V483A | 28 | 0.2 | S1 domain B (RBD) |
| Q675H | 24 | 0.2 | S1 domain D |
| S50L | 24 | 0.2 | S1 domain A |
| S943P | 22 | 0.1 | S2 HR1 |
| A1078S | 21 | 0.1 | S2 |
| R21I | 19 | 0.1 | S1 domain A |
| V367F | 18 | 0.1 | S1 domain B (RBD) |
| T29I | 18 | 0.1 | S1 domain A |

List of SARS-CoV-2 spike mutations with a frequency ≥ 0.1% in a set of 15335 isolates downloaded from the Global Initiative for Sharing All Influenza Data (GISAID) SARS-CoV-2 sequence database on 5/3/20 (Elbe and Buckland-Merrett, 2017; Shu and McCauley, 2017). The genomes were processed with the nextstrain augur pipeline (https://github.com/nextstrain/augur) (Hadfield et al., 2018), using MAFFT v7.464 (Katoh and Standley, 2013) for sequence alignment and FastTree (Price et al., 2010) to generate a phylogenetic tree. The resulting data were then analyzed with a custom Swift program.

**Figure S1 (related to Figures 2 and 3)**

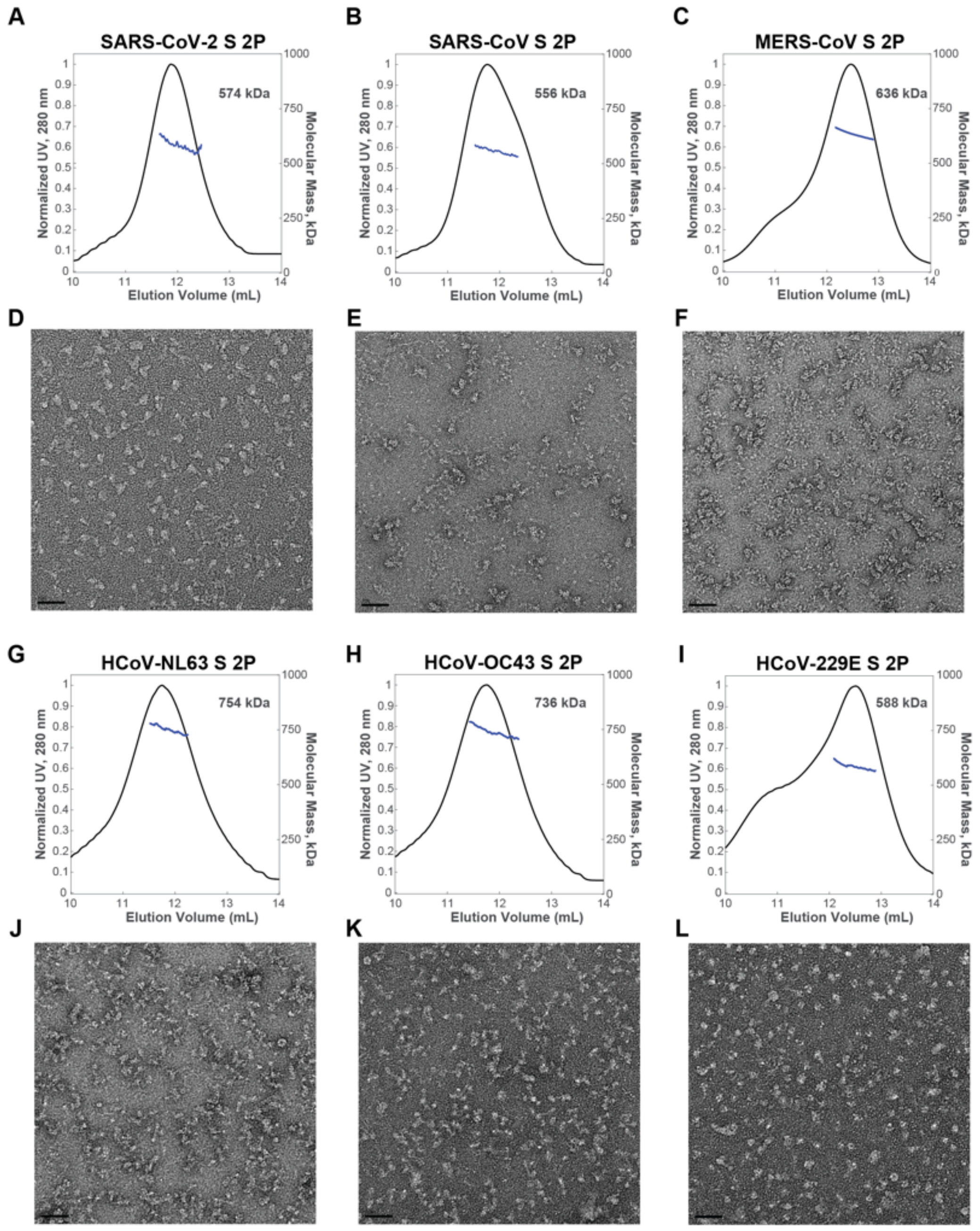

**Figure S1. SEC-MALS characterization of purified proteins (related to Figures 2 and 3).**

(A-C; G-I) SEC-MALS of CoV S trimers and (D-F; J-L) corresponding representative nsEM. Scale bar on micrographs represents 50 nm.

**Figure S2 (related to Figure 3)**

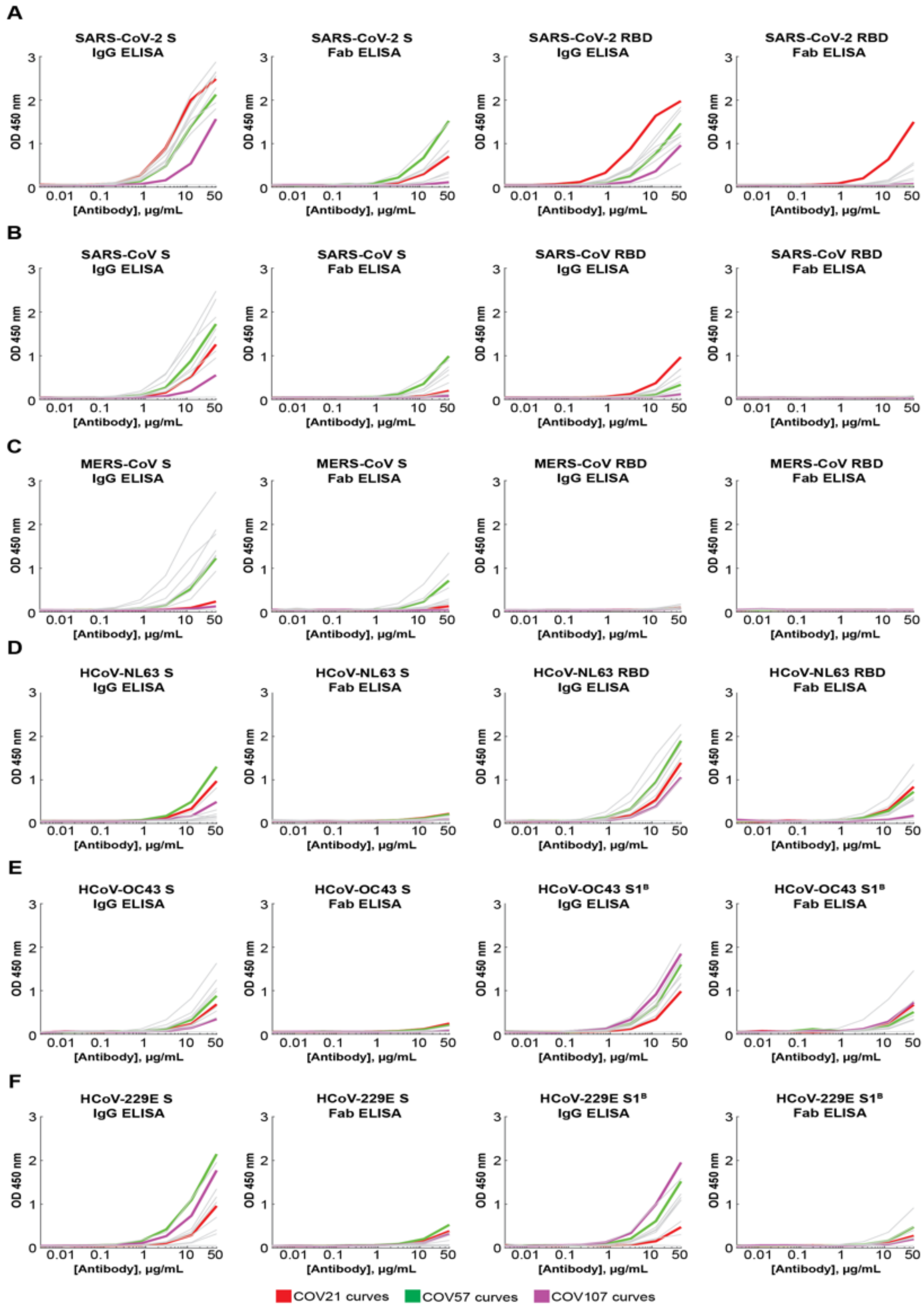

**Figure S2. SARS-CoV-2, SARS-CoV, MERS-CoV and common cold coronavirus ELISA curves (related to Figure 3)**. Anti-S IgG (left panel), Anti-S Fab (middle left panel), Anti-RBD/S1$^{B}$ IgG (middle right panel), and Anti-RBD/S1$^{B}$ Fab (right panel) ELISA binding data for (A) SARS-CoV-2, (B) SARS-CoV, (C) MERS-CoV, (D) HCoV-NL63, (E) HCoV-OC43, and (F) HCoV-229E. COV21: red curves; COV57: green curves; COV107: magenta curves. Curves for other plasmas are in gray. Each curve represents the average of three independent experiments. Binding of the IgG and Fab from IOMA, an antibody against HIV-1 (Gristick et al., 2016), was used as a control in each assay.

### Figure S3 (related to Figure 3)

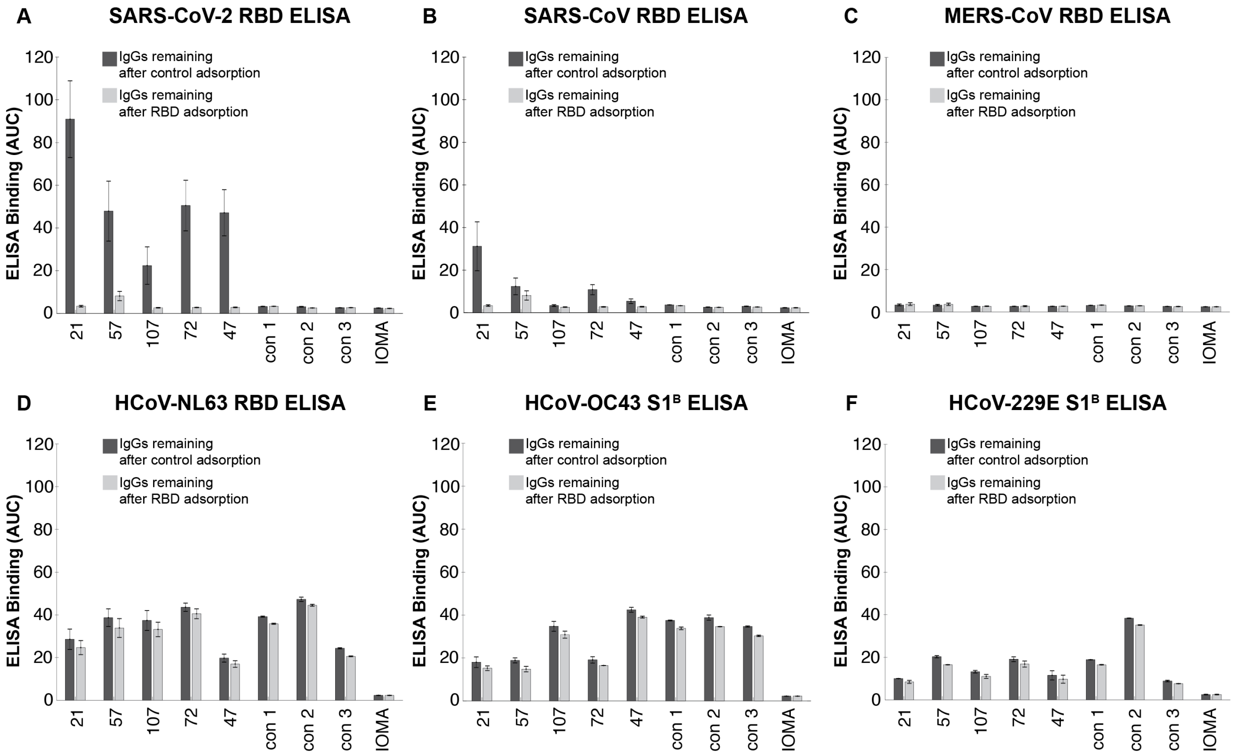

**Figure S3. RBD adsorption experiments to assess degrees of cross-reactive RBD recognition by plasma IgGs (related to Figure 3).** (A-F) Purified IgGs from COVID-19 plasmas (indicated by numbers) and control plasmas (indicated as "con") were adsorbed with one of two resins: a SARS-CoV-2 RBD resin (IgGs remaining after RBD adsorption; light gray bars) and a 2G12 mAb control resin (IgGs remaining after control adsorption; dark gray bars). IgGs remaining after adsorption were evaluated in ELISAs against the indicated RBD (or $S1^B$) domains. Binding of IgGs after adsorption to IOMA, an antibody against HIV-1 (Gristick et al., 2016), was used as a control in each assay. Results are presented as area under the curve (AUC; shown as mean of experiments conducted in duplicate).

**Figure S4 (related to Figure 4)**

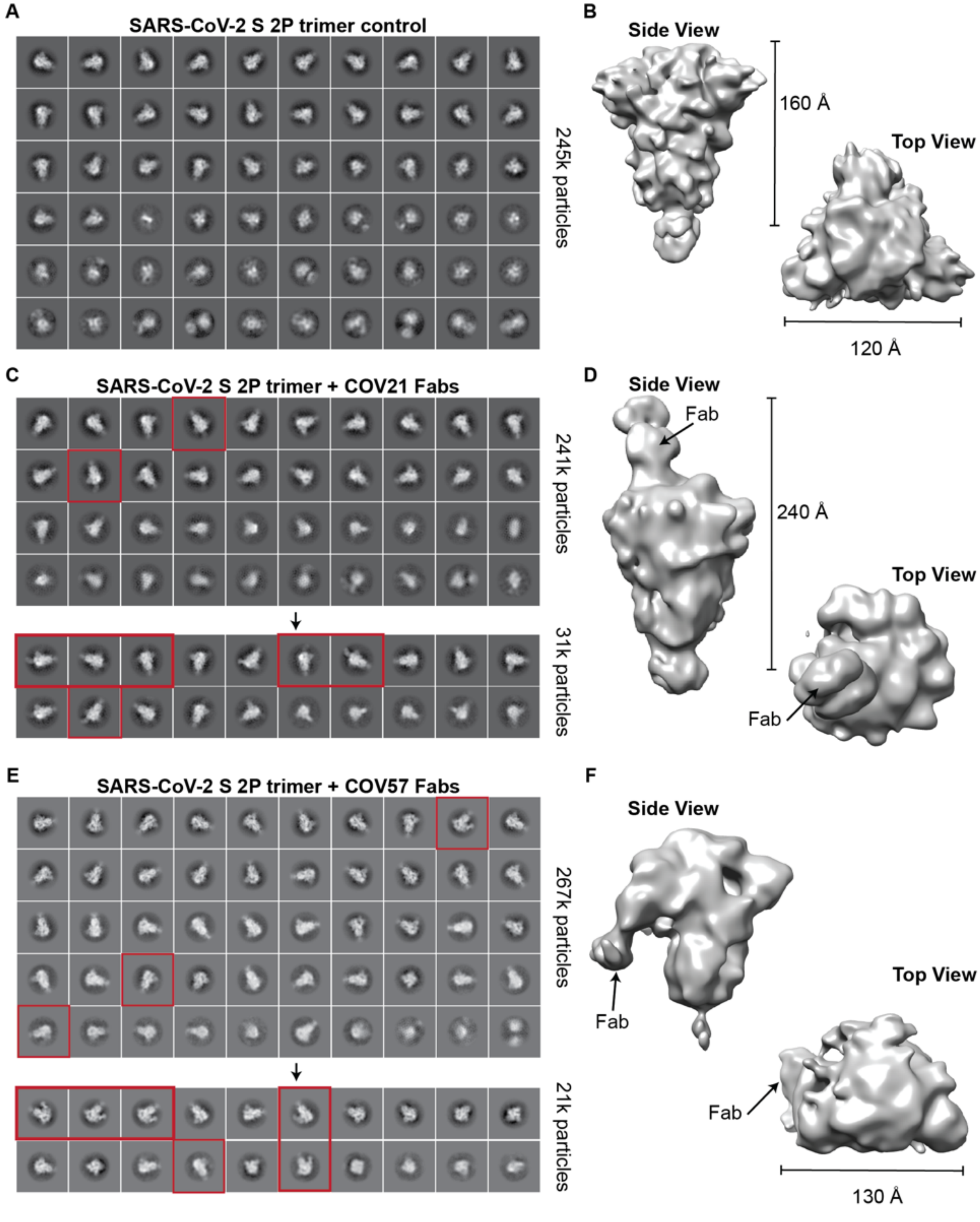

**Figure S4. Representative 2D class-averages and 3D models from nsEMPEM of human convalescent plasma (related to Figure 4)**. (A,C,E) Representative reference-free 2D class-averages obtained from EM data collections of (A) SARS-CoV-2 S trimers alone, (C) SARS-CoV-2 S complexed with COV21 polyclonal Fabs, and (E) SARS-CoV-2 S complexed with COV57 polyclonal Fabs. For COV21 and COV57, class-averages demonstrating extra density beyond the S trimer core are highlighted (red boxes). For COV107, no extra density was observed in class averages or a 3D construction (data not shown). (B,D,F) Refined 3D models after iterative rounds of 2D and 3D classification. Features corresponding to Fabs are denoted.

**Figure S5 (related to Figure 5)**

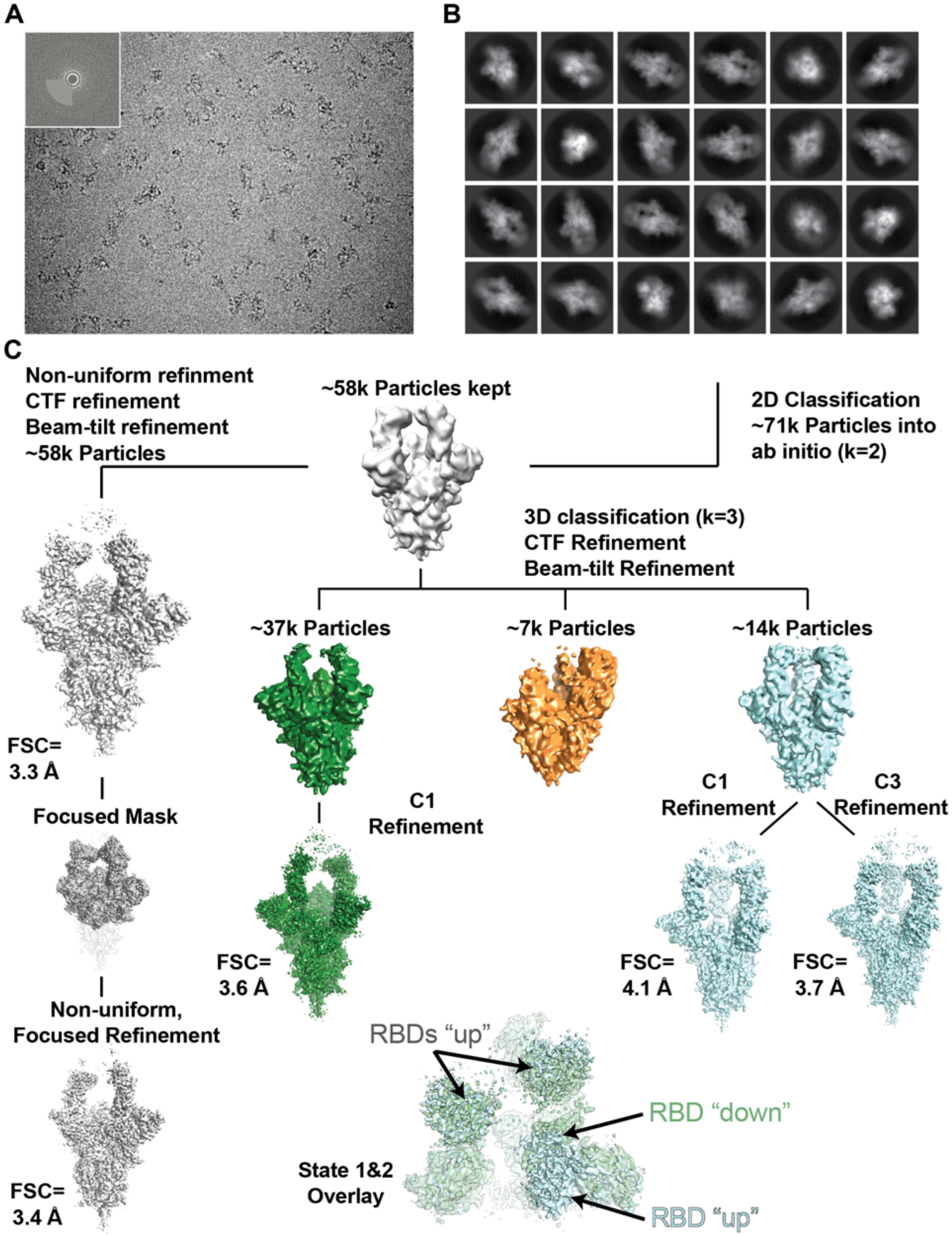

**Figure S5. Data collection and processing pipeline for the cryo-EM structure of the C105-SARS-CoV-2 S complex (related to Figure 5)**. (A) Representative micrograph of C105-S complex in vitreous ice. Power spectrum of micrograph determined during CTF estimation showing Thon rings to 3.2 Å is shown in inset. (B) Reference-free 2D classification of extracted particles. (C) Workflow for classification and refinement of selected particles. Briefly, after selection of good 2D class averages, an ab initio model was generated, which was then homogeneously refined before further 3D classification. To improve features at the SARS-CoV-2 RBD-C105 Fab interface, particles from states 1 and 2 were combined and used for non-uniform, focused refinement to yield a state 1-like reconstruction to an FSC=0.143 resolution of 3.4 Å.

**Figure S6 (related to Figure 5)**

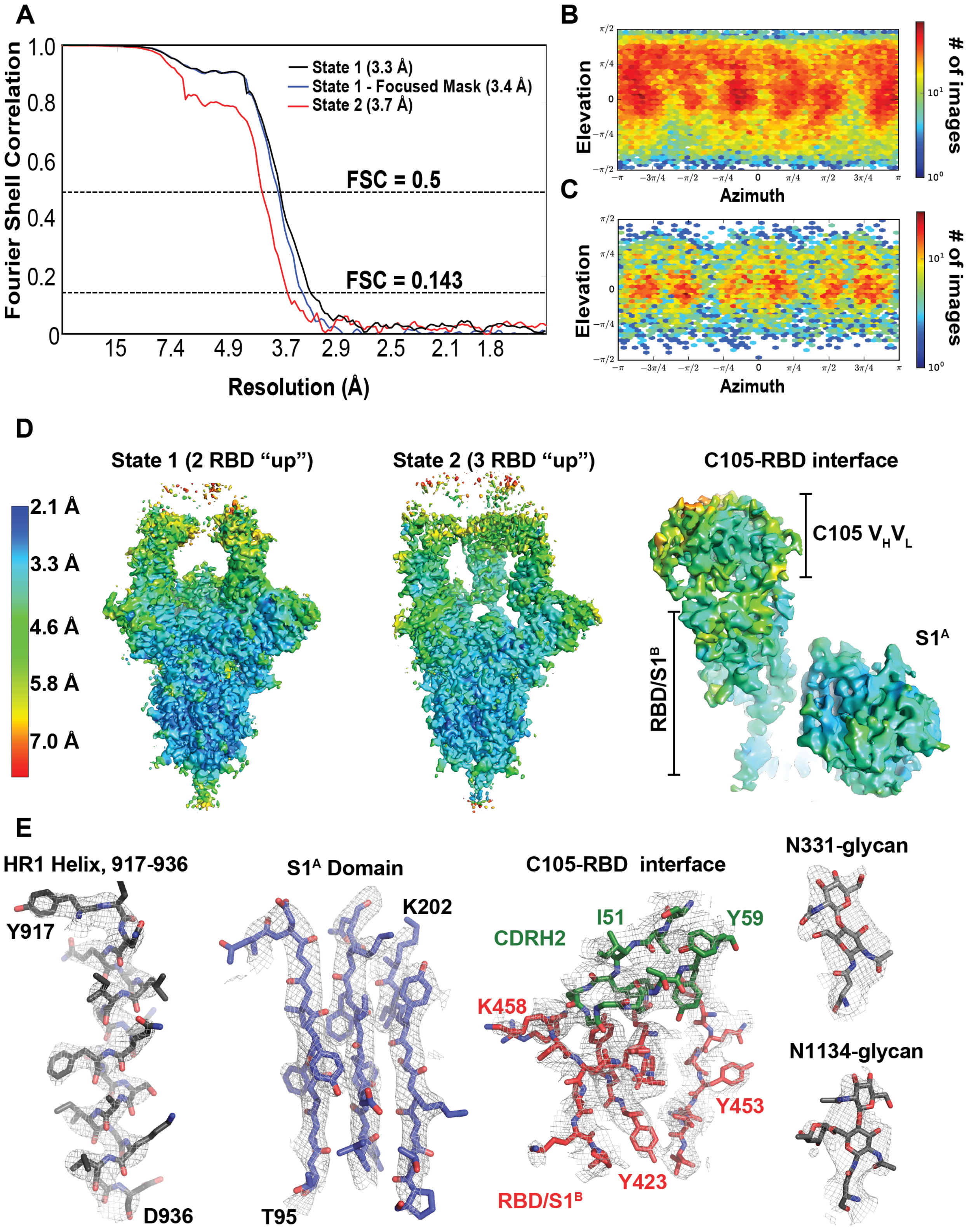

**Figure S6. Cryo-EM structure validation (related to Figure 5)**. (A) Fourier shell correlation (FSC) plots calculated from half-maps of state 1 (black), state 1 after focused refinement (blue) and state 2 (red). Dotted lines for FSC values of 0.5 and 0.143 are shown. (B,C) 2D angular distribution plot for state 1 (panel B) and state 2 (panel C) reconstructions. (D) Local resolution estimations for states 1 and 2 and at the RBD-C105 Fab interface. (E) Representative density from S trimer and Fab regions of the state 1 reconstructed volume. Maps are contoured at 6$\sigma$.

**Figure S7 (related to Figure 5)**

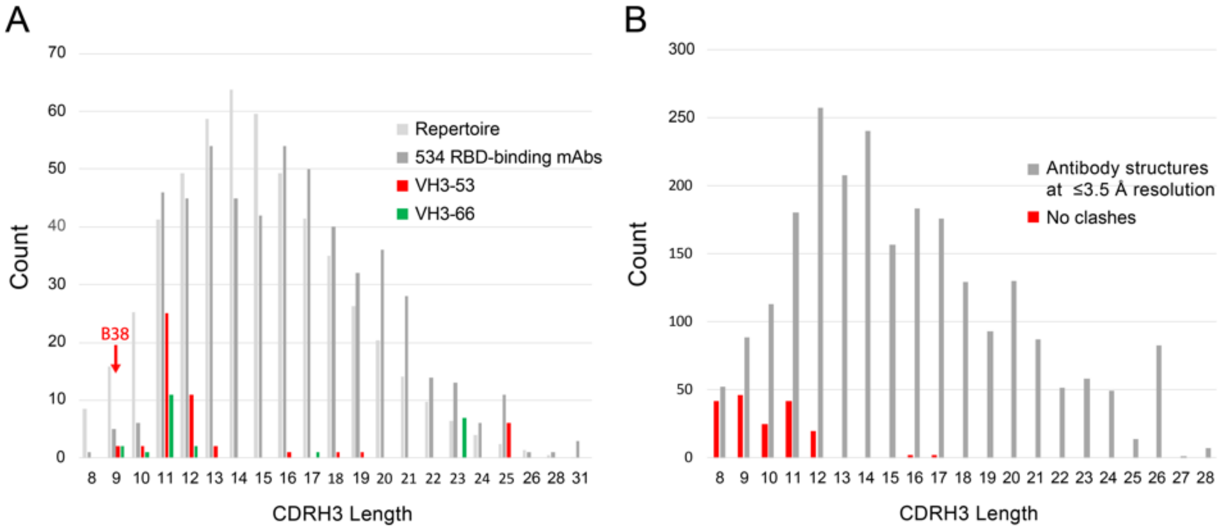

**Figure S7. CDRH3 length distributions (related to Figure 5)**. (A) The CDRH3 lengths (IMGT definition) (Lefranc et al., 2015) of anti-SARS-CoV-2 RBD-binding mAbs (Robbiani et al., 2020) are shown in three groups: all 534 mAbs (dark gray), those derived from *VH3-53* (red), and those derived from *VH3-66* (green). For comparison, the CDRH3 length distribution from the human antibody repertoire (Briney et al., 2019) is also shown (normalized to the same total count as the set of 534). The CDRH3 length of mAb B38 is indicated with an arrow. (B) Length of CDRH3s in human antibodies versus predicted clashes with SARS-CoV-2 RBD if binding in the orientation observed for the mAbs B38 and C105. The VH domains of 1364 human antibody structures with resolutions of ≤3.5 Å downloaded from SAbDab (Dunbar et al., 2014) were aligned to the B38 VH domain in complex with SARS-CoV-2 RBD (PDB code 7BZ5) (Wu et al., 2020c). In cases in which there was more than one Fab in the crystallographic asymmetric unit, each VH was evaluated and enumerated separately. CDRH3 clashes were defined if any CDRH3 atom was within 2.0 Å of an atom in the RBD, a stringent criterion devised to account for not allowing CDR flexibility or different side chain rotamer conformations.
